# Regulation of drought tolerance in *Arabidopsis* involves PLATZ4-mediated transcriptional suppression of *PIP2;8*

**DOI:** 10.1101/2021.10.06.463369

**Authors:** Miao Liu, Chunyan Wang, Zhen Ji, Lei Zhang, Chunlong Li, Jinguang Huang, Guodong Yang, Kang Yan, Shizhong Zhang, Chengchao Zheng, Changai Wu

## Abstract

PLATZ transcription factors play important roles in plant growth, development, and biotic and abiotic stress responses. However, how PLATZ regulates plant drought tolerance and ABA sensitivity remains largely unknown. Here, we show that PLATZ4 increases drought tolerance and ABA sensitivity in *Arabidopsis thaliana* by suppressing the expression of *PIP2;8, DEG3*, At1g33840 and At3g01345, while upregulating the expression of *PR1, ABI3, ABI4* and *ABI5*. Moreover, PLATZ4 directly binds A/T-rich sequences within the *PIP2;8* promoter but not *ABI3, ABI4* and *ABI5* promoters. Consistent with this, *PIP2;8* acts epistatically to *PLATZ4*. Furthermore, the aquaporin activity of PIP2;8 was confirmed in *Xenopus laevis oocytes* and yeast cells in response to osmotic stress. Analysis of water loss of seedlings overexpressing *PIP2;8* or lacking *PIP2;8* function indicated that PIP2;8-mediated water flow is particularly active in response to drought stress *in planta*. Accordingly, stomatal closure was enhanced in *pip2;8* mutants but reduced in *PIP2;8*-overexpressing plants. In *platz4* mutant and *PLATZ*4-overexpressing plants, water loss and stomatal closure changed oppositely to those in *pip2;8* mutants and *PIP2;8-*overexpressing plants, respectively. In addition, the interaction between PLATZ4 and AITR6 was confirmed by several assays, and the binding of *PIP2;8* promoter by PLATZ4 was strengthened by an interaction with AITR6. Collectively, our findings reveal that PLATZ4 interacts with AITR6 to increase ABA sensitivity and drought tolerance in *Arabidopsis* by upregulating expression of *PR1, ABI3, ABI4* and *ABI5* while inhibiting the expression of *PIP2;8* and associated genes.

**One sentence summary:** PLATZ4 interacted with AITR6 to increase drought tolerance and ABA sensitivity in *Arabidopsis thaliana* by directly bind A/T-rich sequences within the plasma membrane aquaporin (PIP)*2;8* promoter.

## Introduction

PLATZ transcription factors (TFs) are plant-specific zinc-dependent DNA-binding proteins. Accumulating evidence has revealed the important roles played by PLATZs in many processes related to plant growth, development, and responses to environmental stimuli, including stem cell niche and root meristem size regulation(Yamada et al., 2020), endosperm development and seed filling(Li et al., 2017; Wang et al., 2019a), leaf cell proliferation and leaf senescence(Kim et al., 2018), desiccation tolerance in seeds, drought tolerance in vegetative tissues(González-Morales et al., 2016), and response to salt and osmotic stresses(Liu et al., 2020; Zhang et al., 2018). In previous work, we showed that PLATZ4 (At1g32700), named in this study, is significantly induced by mannitol and interacted with MYBs and ABA-induced transcription repressor 6 (AITR6) in *Arabidopsis*, except for PLATZ2 (Liu *et al*., 2020). The function of AITR6 as a transcriptional repressor in regulating ABA and drought stress response indicates a possible role of PLATZ4 in ABA and drought stress regulation in *Arabidopsis*(Song et al., 2016; Tian et al., 2017). However, no direct evidence has been provided.

Drought is a major abiotic stress that affects agricultural production worldwide, and increasing research effort is being devoted to development of drought-tolerant crop plants. Aquaporins (AQPs) are primarily involved in maintaining cellular water homeostasis(Wang et al., 2020; Zargar et al., 2017). Several lines of evidence indicate that AQPs are responsible for precise regulation of water movement and consequently play a crucial role in drought tolerance(Zargar *et al*., 2017). Thus, AQPs are key candidate molecules for the development of drought-tolerant crops. Hence, there is an urgent need to explore the potential AQPs by understanding the molecular mechanisms and pathways by which they induce drought tolerance.

The diverse plant AQP protein family is classified into five subfamilies: plasma membrane intrinsic proteins (PIPs), tonoplast intrinsic proteins (TIPs), NOD26-like intrinsic proteins (NIPs), small basic intrinsic proteins (SIPs), and unclassified X intrinsic proteins (XIPs)(Bienert and Bienert, 2017). The PIPs and TIPs are predominantly involved in water transport(Deshmukh et al., 2017; Wang *et al*., 2020), and have consequently been the most extensively studied, particularly under water stress conditions. Multiple environmental factors, such as drought, salinity, cold, and light, regulate AQPs, mostly at the transcriptional level(Liu et al., 2013a). However, only a few transcription factors have been shown to be directly involved in the regulation of *AQP* gene expression. MdDREB6.2 binds the promoters of *MdPIP1;3* and *Mdγ-TIP* to active their expression, and thus contributes to the drought stress response(Wang et al., 2017). ABI3 binds directly to the RY motif of the *TIP3* promoter to maintain seed longevity during seed maturation(Mao and Sun, 2015). ASR1 from tomato binds the GCCCA motif in the promoter of *PIP1;4* to induce its expression in response to drought stress(Ricardi et al., 2014). Hence, our knowledge of the direct transcriptional regulators of *AQP* genes remains limited.

In this study, we revealed that PLATZ4 positively regulated ABA response and drought tolerance in *Arabidopsis*. Further investigations indicated that PLATZ4 regulates the expression of key genes involved in ABA signaling and drought tolerance, including *ABI3, ABI4, ABI5, PIP2;8*, and *PR1*. Direct binding of PLATZ4 to the *PIP2;8* promoter was verified by yeast one hybrid (Y1H), chromatin immunoprecipitation followed by quantitative PCR (ChIP-qPCR), and electrophoretic mobility shift assay (EMSA) experiments. Under drought stress treatment, *Arabidopsis platz4* mutants exhibited reduced tolerance to drought stress, whereas the *pip2;8* mutant exhibited higher drought tolerance than the wild type (WT). Collectively, our findings reveal a novel PLATZ transcription factor that directly inhibits the expression of *PIP2;8* to increase drought tolerance in *Arabidopsis*.

## Results

### PLATZ4 enhances drought tolerance in *Arabidopsis*

To characterize the function of *PLAZT4* in *Arbidopsis*, we generated *PLAZT4* overexpression (P4OE) and RNA-interference (P4RNAi) lines. The transcripts of *PLAZT4* in P4OE and P4RNAi seedlings were detected by RT-PCR (Figure 1A and 1B). Three *PLAZT4* overexpression lines (P4OE#2, P4OE#4 and P4OE#5) and two *PLATZ4* RNAi lines (P4RNAi3 and P4RNAi9) were used for phenotypic analysis.

**Figure 1.**
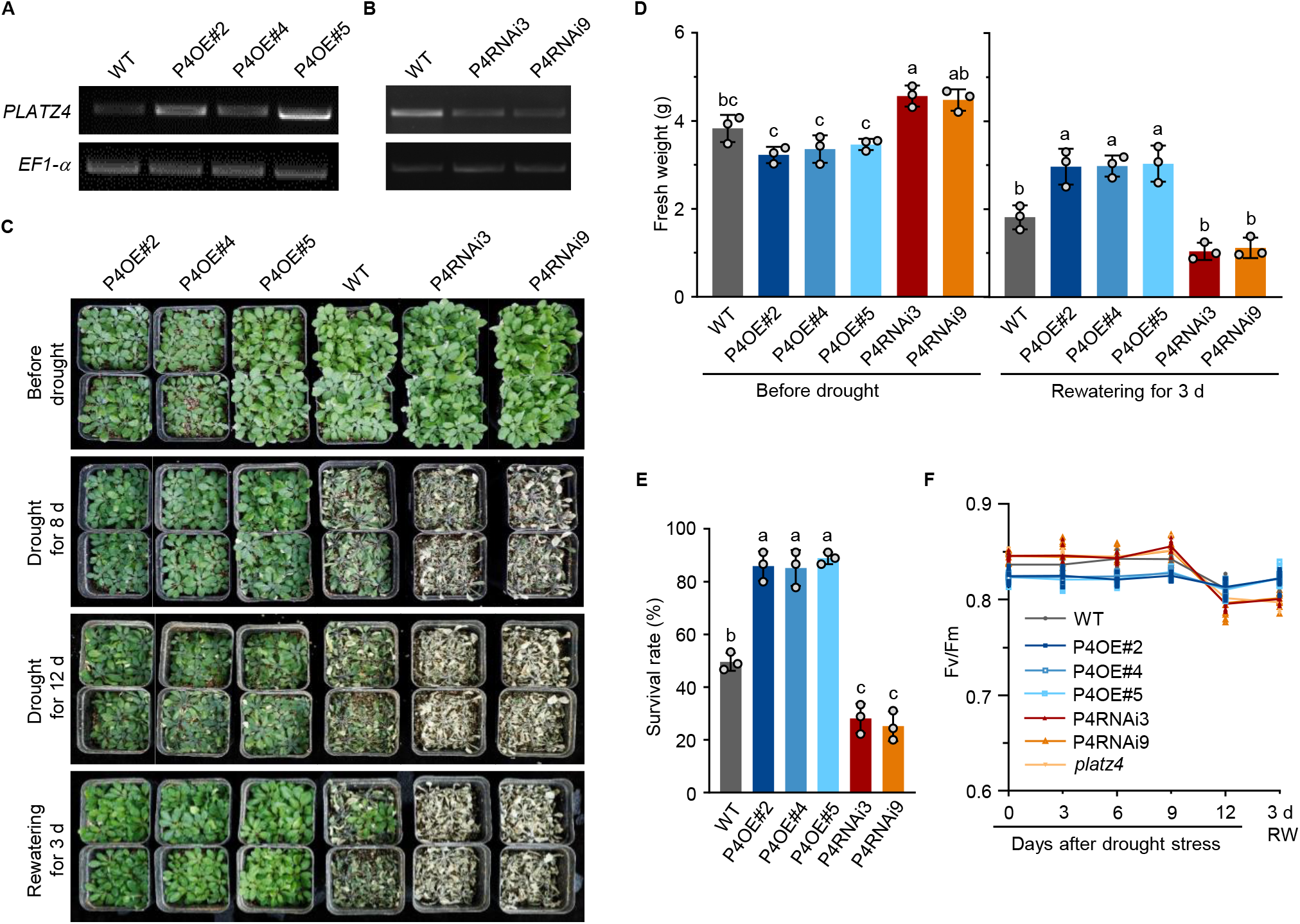
PLATZ4 enhances drought tolerance in *Arabidopsis*. (A,B) Transcript levels of *PLATZ4* in WT, *PLATZ4* overexpressing and RNAi *Arabidopsis* determined by RT–PCR. *EF-1α* was used as the internal control. (C) Drought tolerance assays of WT, P4OE#2, P4OE#4, and P4OE#5 and P4RNAi3, P4RNAi9 plants. Three-week-old plants were subjected to drought stress by no watering for 8 or 12 d and then rewatered for 3 d. (D,E) Fresh weight and survival rates of plants in (C). Survival rates and fresh weight were recorded 3d after rehydration. Error bars indicate SEM (*n* = 3). Bars labeled with different lowercase letters are significantly different from one another (*P* < 0.05; one-way ANOVA). (F) Effects of drought stress on Fv/Fm values in leaves of WT, P4OE#2, P4OE#4, P4OE#5, and P4RNAi3, *platz4* plants. 15-day-old plants were subjected to drought stress by no watering for 0, 3, 6, 9, and 12 d, and then rewatered (RW) for 3 d. F_0_ and Fm were tested at the indicated time points. Fv/Fm was calculated by (Fm-F_0_)/Fm. Values represent means ± SD (*n* >10) from three biological replicates.

Before drought stress treatment (Figure 1C), P4OE seedlings were smaller and P4RNAi seedlings were larger than WT. After drought for 8 d, P4OE seedlings grew well, whereas P4RNAi plants died. After drought for 12 d, P4OE seedlings exhibited serious wilt and most WT seedlings died (Figure 1C). And after rewatering for 3 d, most P4OE seedlings recovered (survival rate, 82%), whereas 50% of WT and 30% of P4RNAi seedlings survived (Figure 1E). Fresh weight was higher than WT in P4OE seedlings and lower than WT in P4RNAi (Figure 1D). To further confirm the function of *PLATZ4* in P4RNAi seedlings, we generated a T-DNA insertion mutant of *PLATZ4* and verified it by RT-PCR (Supplemental Figure 1A and 1B). Phenotype, survival rate, and fresh weight indicated that the *platz4* mutant was less drought tolerant than WT (Supplemental Figure 1). The *platz4* phenotype was similar to those of P4RNAi lines, indicating that *PLATZ4*, but not other *PLATZ*s, functions in P4RNAi seedlings. Consistent with this, under normal and early drought stress, the Fv/Fm ratios of P4OE leaves were lower than those of WT, whereas those of P4RNAi and *platz4* seedlings were higher. However, under severe drought stress, the Fv/Fm values of P4OE leaves decreased slightly, whereas those of P4RNAi and *platz4* leaves decreased sharply and were lower than that of P4OE leaves (Figure 1F). Taken together, these results indicate that PLATZ4 positively regulates drought tolerance in *Arabidopsis*.

### PLATZ4 enhances ABA sensitiviy in *Arabidopsis*

In general, ABA mediates plant drought tolerance. Therefore, we analyzed the ABA response. On half-strength Murashige and Skoog (MS) medium without ABA, the fresh weight of the P4OE seedlings was 10–20% lower during germination stage (Figure 2B), indicating *PLAZT4* overexpression lines germinated and grew more slowly than WT. In the presence of 1 μM ABA, the P4OE seedlings grew significantly slower than WT, with 60–70% lower fresh weight and about 45% shorter root length (Figure 2A and 2C). In the presence of 2 μM ABA, the growth of P4OE seedlings was further inhibited, and the roots were shorter. By contrast, P4RNAi and *platz4* seedlings had higher fresh weight and longer roots than WT under both ABA conditions (Figure 2D-2F). When 4-day-old uniformly developed seedlings were transferred onto half-strength MS medium with 5 or 10 μM ABA, the root lengths of the P4OE seedlings were shorter than WT, whereas those of P4RNAi3 and *platz4* seedlings were longer than WT (Supplemental Figure 2C and 2D). These data indicate a positive role of PLATZ4 in the ABA response during seed germination and seedling growth.

**Figure 2.**
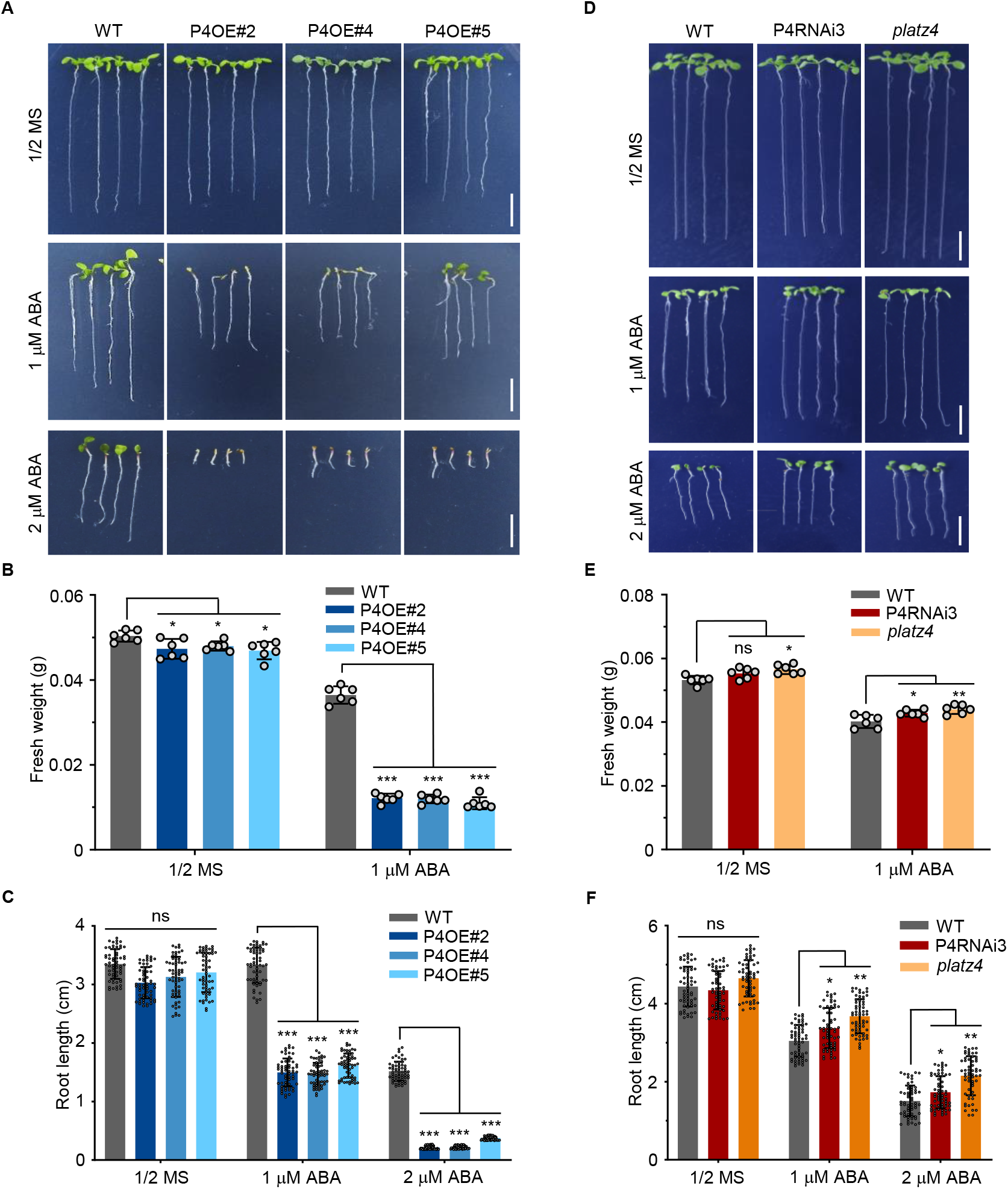
PLATZ4 enhances ABA sensitivity in *Arabidopsis*. (A) Phenotypes of WT, P4OE#2, P4OE#4, and P4OE#5 seedlings grown for 8 d on halfstrength MS medium with or without 1 or 2 μM ABA. (B) Fresh weight of the seedlings shown in (A). Error bars indicate SEM (*n* = 6). The asterisks indicate significant differences compared with WT. *, *P* < 0.05; ***, *P* < 0.001 (one-way ANOVA). (C) The primary root length of the seedlings shown in (A). Error bars indicate SEM (*n* = 60). The asterisks indicate significant differences compared with WT. ns, not significant; ***, *P* < 0.001 (one-way ANOVA). (D) Phenotypes of WT, P4RNAi3 and *platz4* seedlings grown for 10 d on half-strength MS medium with or without 1 or 2 mM ABA. (E) Fresh weight of the seedlings shown in (D). Error bars indicate SEM (*n* = 6). The asterisks indicate significant differences compared with WT. ns, not significant; *, *P* < 0.05; **, *P*< 0.01 (one-way ANOVA). (F) The primary root length of the seedlings shown in (D). Error bars indicate SEM (*n* = 60). The asterisks indicate significant differences compared with WT. ns, not significant; *, *P* < 0.05; **, *P* < 0.01 (one-way ANOVA). These experiments were repeated three times with similar results.

To determine whether PLATZ4 affects ABA biosynthesis or signaling, we examined the expression of genes involved in ABA biosynthesis, such as *ABA1, ABA2, ABA3, NECD3*, and *NCED9*, or ABA signaling, such as *ABI1, ABI3, ABI4*, and *ABI5* (Supplemental Figure 3). Reverse transcription–quantitative PCR (RT-qPCR) analysis revealed that the expression of *ABA1, ABA2, ABA3, NECD3, NCED9*, and *ABI1* was not changed in transgenic seedlings relative to WT, whereas the expression of *ABI3, ABI4*, and *ABI5* was significantly up-regulated in the P4OE seedlings, but not in P4RNAi or *platz4* seedlings. Next, we sought to determine whether PLATZ4 directly targets the promoters of *ABI3, ABI4*, and *ABI5*. To this end, we first performed Y1H experiments. Three DNA fragments from each of the *ABI3, ABI4*, and *ABI5* promoters were cloned into pLacZi2u vector (Supplemental Figure 4). Simultaneously, PLATZ4 was cloned into pJG4-5 vector. The Y1H results revealed that PLATZ4 did not bind any of the DNA fragments within the *ABI3, ABI4*, and *ABI5* promoters (Supplemental Figure 4B).

Next, we performed a Dual-Luciferase Reporter Assay. The same DNA fragments from the *ABI3, ABI4*, and *ABI5* promoters were introduced into a vector to generate the reporter constructs, and PLATZ4 driven by the 35S promoter was used as the effector. *Pro35S:GFP* was used as a control (Supplemental Figure 4C). For all of the DNA fragments, the ratio of LUC/RenLUC did not differ significantly between PLATZ4 and the GFP control (Supplemental Figure 4D), indicating that PLATZ4 did not bind the *ABI3, ABI4*, or *ABI5* promoters directly. Together, these observations indicate that PLATZ4 indirectly and positively regulates ABA signaling but not ABA biosynthesis.

### PLATZ4 represses *PIP2;8* expression by directly targeting the A/T rich sequences in its promoter

To investigate the molecular mechanism by which PLATZ4 contributes to ABA sensitivity and drought tolerance, we analyzed the transcriptional profiles of P4OE#2 and WT plants. The results revealed that PLATZ4 significantly regulated 505 and 477 differently expressed genes (DEGs), out of 23144 genes analyzed under normal and osmotic conditions, respectively (Supplementary Table 1 and Supplemental Figure 5). Criteria for significant regulation were |logFC| ≥1 and p value < 0.05. Gene ontology analysis showed that the genes differentially regulated under normal conditions were mainly involved in defense, stress, chemical responses, auxin signaling, and the transport of nitrate, ions, and water (Supplemental Figure 6A). The genes differentially regulated after 3 h of drought stress were mainly involved in transcription factor activity, oxidoreductase activity, cell death, and transport of anions, potassium, sodium, and water (Supplemental Figure 6B). Some DEGs involved in abiotic stress responses were verified by RT-qPCR (Supplemental Figure 7A). The results showed that *PIP2;8* (At2g16850), *PLATZ12* (At5g46710), *DEG3*, At3g01345, and At1g33840 were significantly downregulated, whereas *CYSTM8* and *PR1* were obviously upregulated by *PLATZ4* overexpression. The results of RT-qPCR revealed a high correlation with the transcriptional profiles (R^2^=0.847) (Supplemental Figure 7B), confirming the high reliability of the transcriptional profiles.

One of the DEGs, *PIP2;8*, encodes a plasma membrane aquaporin for water transport, and we speculated that it was a target of PLATZ4. To test this hypothesis, we monitored expression of *PIP2;8* in P4OE, P4RNAi3, and *platz4* seedlings. *PIP2;8* expression was significantly reduced relative to WT in the P4OE lines, but upregulated in P4RNAi3 and *platz4* plants under both normal and osmotic conditions (Figure 3A). Next, we assessed the effect of PLATZ4 on *PIP2;8* promoter activity by detecting the GUS activity under the control of the *PIP2;8* promoter (−1~−1813 bp) (*ProPIP2;8:GUS*) in *Nicotiana benthamiana* leaves. PLATZ4 repressed GUS activity to about 50% of control levels (Supplemental Figure 8A), supporting a role for PLATZ4 in repressing *PIP2;8* expression. For this purpose, we performed Y1H experiments. Three DNA fragments from the *PIP2;8* promoter were cloned into pLacZi2u (Figure 3B). The Y1H results revealed that PLATZ4 strongly bound the F3 region (bp −930 – −1) within the *PIP2;8* promoter (Figure 3B). Next, we performed a Dual-Luciferase Reporter Assay. Six fragments including F4–F9 within the F3 region of the *PIP2;8* promoter were introduced into a vector to generate reporter constructs; *Pro35S:PLATZ4* was used as the effector, and *Pro35S:GFP* was used as a control (Supplemental Figure 8B). The ratio of LUC/RenLUC reported by the F3–F5 DNA fragments were lowered by PLATZ4 compared with the GFP control, while that by F6-F9 DNA fragments were not changed (Figure 3C), indicating that PLATZ4 bound the region from −930 to −626 bp within the *PIP2;8* promoter (Supplemental Figure 8B). ChIP-qPCR was performed to amplify the F10–F14 DNA fragments within the *PIP2;8* promoter and only the F12 (−880 – −708 bp) DNA fragment was associated with PLATZ4-HA (Supplemental Figure 8C and Figure 3D). These results illustrate that PLATZ4 directly binds the −880 – −708 bp region of the *PIP2;8* promoter.

**Figure 3.**
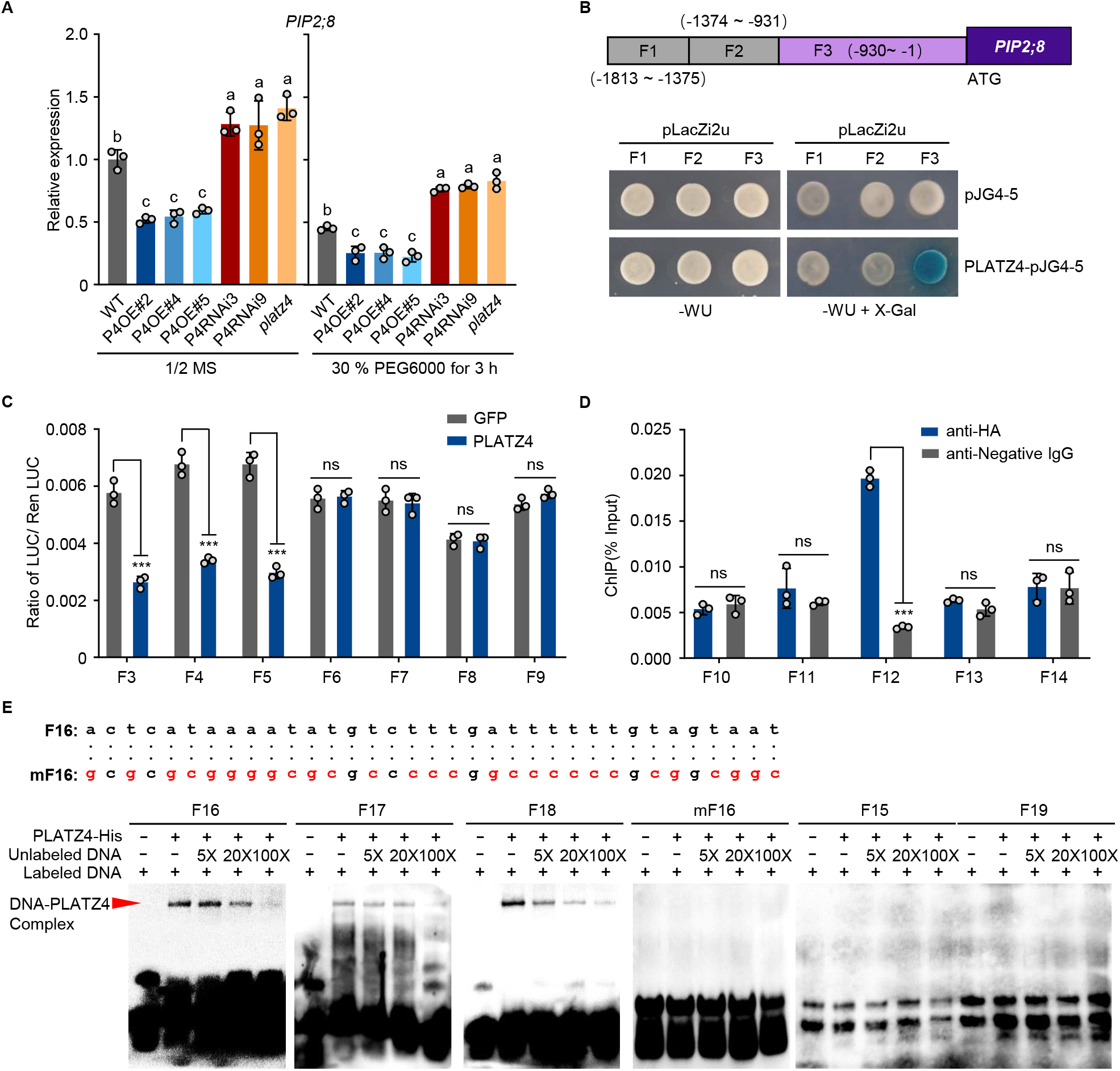
PLATZ4 represses *PIP2;8* expression by directly targeting the A/T rich sequences in its promoter. (A) Transcript levels of *PIP2;8* in 15-day-old seedlings of WT, P4OE#2, P4OE#4, P4OE#5, P4RNAi3, P4RNAi9 and *platz4* that were grown under half-strength MS with 30% PEG6000 for 3 h or not. Error bars indicate SEM (*n* = 3). Bars labeled with different lowercase letters are significantly different from one another (*P* < 0.05; one-way ANOVA). (B) Yeast one-hybrid (Y1H) assays revealed the binding of PLATZ4 to the F3 DNA fragment in *PIP2;8* promoter. F1 to F3 indicate the three DNA fragments within *PIP2;8* promoter. Yeast cells were co-transformed with GAL4-Activation Domain (pJG4-5) or PLATZ fused with GAD (PLATZ4-pJG4-5), and the LacZ reporter gene was driven by F1 to F3 DNA fragments (*ProF1:LacZ, ProF2:LacZ, ProF3:LacZ*) and grown on medium containing X-Gal. Co-expression of GAD and *ProF1:LacZ, ProF2:LacZ*, or *ProF3:LacZ* was used as a negative control. WU, Synthetic Dropout/-Trp-Ura. (C) In vivo transient Dual-Luciferase Reporter Assays verified the binding of PLATZ4 to the F3 to F5 DNA fragments in *PIP2;8* promoter in *N. benthamiana* leaves. Effector constructs contained the PLATZ4 driven by the CaMV35S promoter. Reporter constructs contained the indicated DNA fragments within *PIP2;8* promoter. And *Pro35S:GFP* was used as the negative control. The ratios of LUC activities to REN activities (LUC/REN) were means ± SD (*n* = 3). ns, not significant; ***, *P* < 0.001 (Student’s t-test). (D) ChIP-qPCR analysis to determine the PLATZ4 binding regions in the *PIP2;8* promoter. Chromatin was isolated from 7-day-old seedlings of the *ProSuper:PLATZ4-HA* transgenic *Arabidopsis* grown on half-strength MS medium. Immunoprecipitation was carried out with the HA antibody. Then the five DNA fragments in the *PIP2;8* promoter were amplified by qPCR using sequence specific primers. Error bars indicate SEM (*n* = 3). ns, not significant; ***, *P* < 0.001 (Student’s t-test). (E) EMSA assays show the binding sites of PLATZ4 in the *PIP2;8* promoter. The sequences of F15 to F19 DNA fragments in the *PIP2;8* promoter were listed in Supplementary Table S2. The F16 DNA fragment was mutant to mF16 DNA fragment as shown in (E). Arrows indicate the shifted band of the protein–DNA complex. The plus (+) and minus (−) signs denote the presence or absence of the protein or DNA in each sample. An excess of the unlabeled probe was added to compete with the labeled probe.

To further confirm the direct binding of PLATZ4 to the −880 – 708 bp region of *PIP2;8* promoter, we performed EMSAs. A/T-rich sequences have been identified as PLATZ recognition sites in the promoters of PLATZ target genes in pea(Nagano et al., 2001),

*Arabidopsis*(González-Morales *et al*., 2016; Kim *et al*., 2018; Liu *et al*., 2020), Zea mays(Li *et al*., 2017; Wang et al., 2018), and rice(Wang et al., 2019b). Three DNA fragments (F16-F18) containing A/T-rich sequence, two DNA fragments (F15 and F19) not containing A/T-rich sequence, and a mutant F16 DNA fragment (mF16) within the −880 – −708 bp region of the *PIP2;8* promoter were synthesized and labeled by biotin as probes (Supplemental Figure 8D, Figure 3E and Supplementary Table 2). The same unlabeled DNA were used as competitor. As shown in Figure 3E, when the purified PLATZ4-GST was incubated with the labeled F16 probe, a protein-DNA complex with reduced migration was observed, and the binding was inhibited when unlabeled competitor was added, suggesting binding of PLATZ4 to the F16 DNA fragment. Similar results were obtained when purified PLATZ4-GST protein was incubated with labeled F17 and F18 probes with or without the corresponding unlabeled competitors, suggesting that PLATZ4 bound to F17 and F18 DNA fragments. However, when purified PLATZ4-GST protein was incubated with the labeled mF16, F15, and F19 probes with or without the corresponding unlabeled competitors, no protein–DNA complex was detected. These results indicated that PLATZ4 directly bind three DNA sequences within the −827 – −728 bp region of the *PIP2;8* promoter. However, expression of *PIP2;6*, one of the homologous genes of *PIP2;8*, was not changed in P4OE, P4RNAi3 and *platz4* seedlings (Supplemental Figure 9A). Together, the Y1H experiments, Dual-Luciferase Reporter Assays, and ChIP-qPCR results indicated that PLATZ4 did not bind the *PIP2;6* promoter (Supplemental Figure 9B-9E). Taken together, these data demostrate that PLATZ4 specifically targets and suppresses *PIP2;8* expression.

### PIP2;8 functions epistaticaly to *PLATZ4* in regulating drought tolerance in *Arabidopsis*

To test the regulation of PIP2;8 to plant drought tolerance, we generated *PIP2;8* overexpression lines including 2;8OE#2, 2;8OE#38, and 2;8OE#49, as well as *PIP2;8* T-DNA insertion mutants including *pip2;8-1* (Salk_099098) and *pip2;8-2* (SK16840), and verified these lines by RT-PCR (Figure 4A and 4B). These plants were then subjected to drought stress assays. After drought for 12 d and rewatering for 3 d, 2;8OE plants exhibited seriously wilt leaves with significantly lower (20%) survival rates, whereas *pip2;8-1* and *pip2;8-2* plants had full expanded leaves with significantly higher survival rates (68-82%) than the WT (50% survival rate) (Figure 4C and 4D). The fresh weight showed a trend similar to that of survival rate in response to drought stress (Figure 4E), the Fv/Fm ratios of *pip2;8-1* leaves were higher than those of WT, whereas those of 2;8OE leaves were lower than WT (Figure 4F). We also subjected the plants to ABA response assays. In the presence of ABA, the 2;8OE seedlings germinated faster than WT with longer roots; by contrast, *pip2;8-1* and *pip2;8-2* germinated slower with shorter roots (Supplemental Figure 10). These results indicate that *PIP2;8* negatively regulates drought tolerance and ABA sensitivity in *Arabidopsis*.

**Figure 4.**
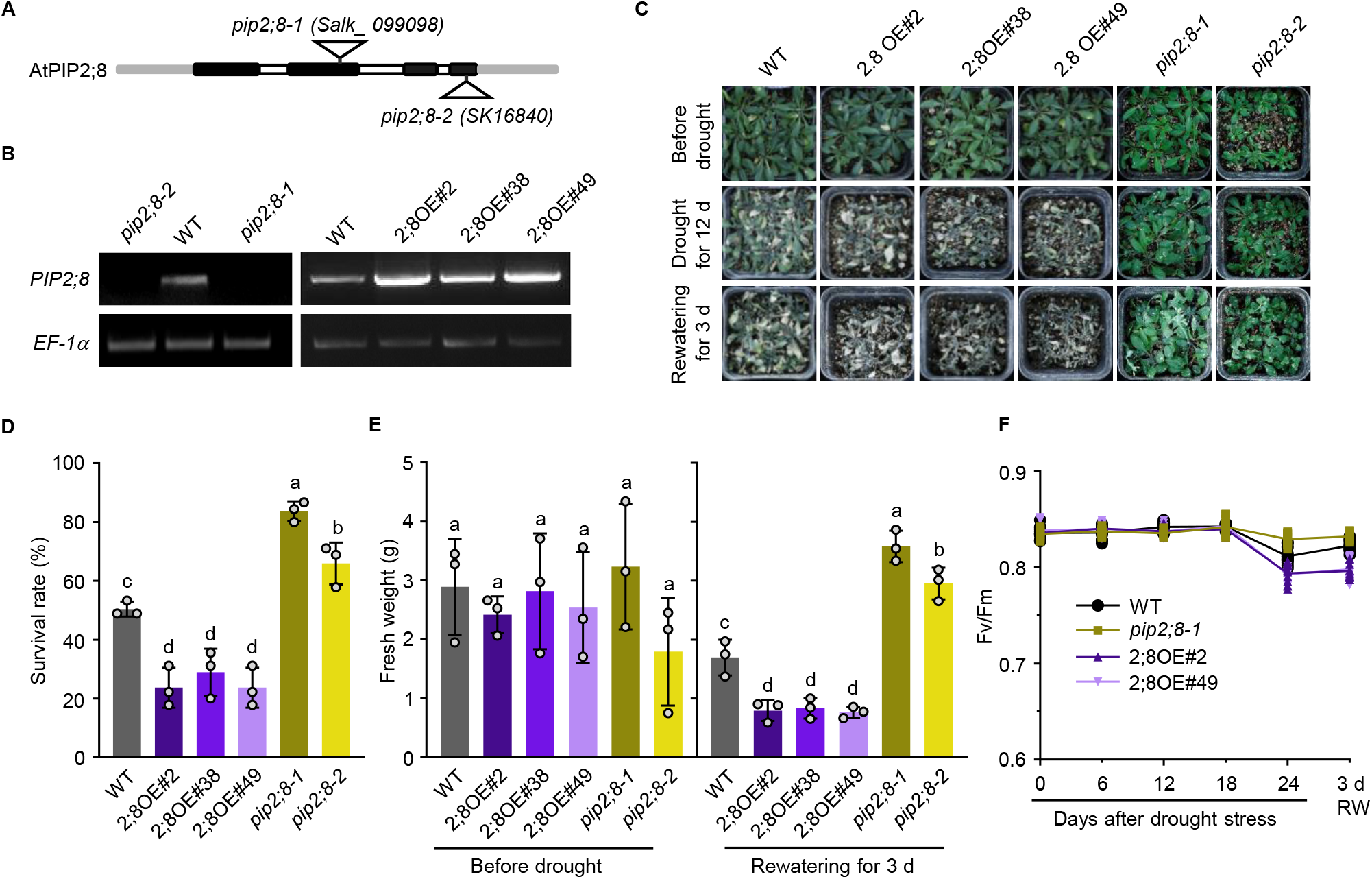
PIP2;8 decreases drought tolerance in *Arabidopsis*. (A) Schematic illustration of the T-DNA insertion sites in the *pip2;8-1* and *pip2;8-2* mutants. (B) Transcript levels of *PIP2;8* in WT, *pip2;8-1* and *pip2;8-2* mutants and transgenic *Arabidopsis* lines overexpressing *PIP2;8* as determined by RT–PCR. *EF-1a* was used as the internal control. (C) Drought tolerance assays of WT, 2;8 OE#2, 2;8 OE#38, 2;8 OE#49 *pip2;8-1*, and *pip2;8-2* plants. Three-week-old seedlings were subjected to drought stress by no watering for 12 d and then rewatering for 3 d. (D,E) Fresh weight and survival rates of seedlings in (C), respectively. Survival were recorded 3 d after rehydration. Fresh weight was measured before drought stress and 3d after rewatering. Error bars indicate SEM (*n* = 3). Bars labeled with different lowercase letters are significantly different from one another (*P* < 0.05; one-way ANOVA). (F) Effects of drought stress on Fv/Fm in leaves of WT, *pip2;8-1*, 2;8OE2, and 2;8OE49 plants. 15-day-old plants were subjected to drought stress no watering for 0, 3, 6, 9, and 12 d, and then rewatered (RW) for 3 d. Fv/Fm was obtained as described in materials and methods. Values represent means ± SD (*n* >10) from three biological replicates.

To confirm that down-regulation of *PIP2;8* is responsible for *PLATZ4* overexpression phenotype, we crossed the 2;8OE#2 and 2;8OE#49 plants with P4OE#2 and P4OE#4 plants to generate four types of *PLATZ4* and *PIP2;8* double-overexpression plants: 2;8OE#2 P4OE#2, 2;8OE#2 P4OE#4, 2;8OE#49 P4OE#2, and 2;8OE#49 P4OE#4. The four types of double-overexpression plants exhibited a drought sensitive phenotype similar to that of 2;8OE plants, which exhibited low survival rates and fresh weight (Figure 5A-5C). In addition, we crossed *pip2;8-1* with P4RNAi3 and *platz4* to generate two types of double mutants,*pip2;8-1* P4RNAi3 and *pip2;8-1 platz4. pip2;8-1* P4RNAi3 and *pip2;8-1 platz4* plants exhibited drought tolerance similar to that of the *pip2;8-1* mutant, which exhibited higher survival rates and fresh weight than WT and P4RNAi3 and *platz4* (Figure 5D-5F). These results demonstrate that *PIP2;8* is epistatic to *PLATZ4*.

**Figure 5.**
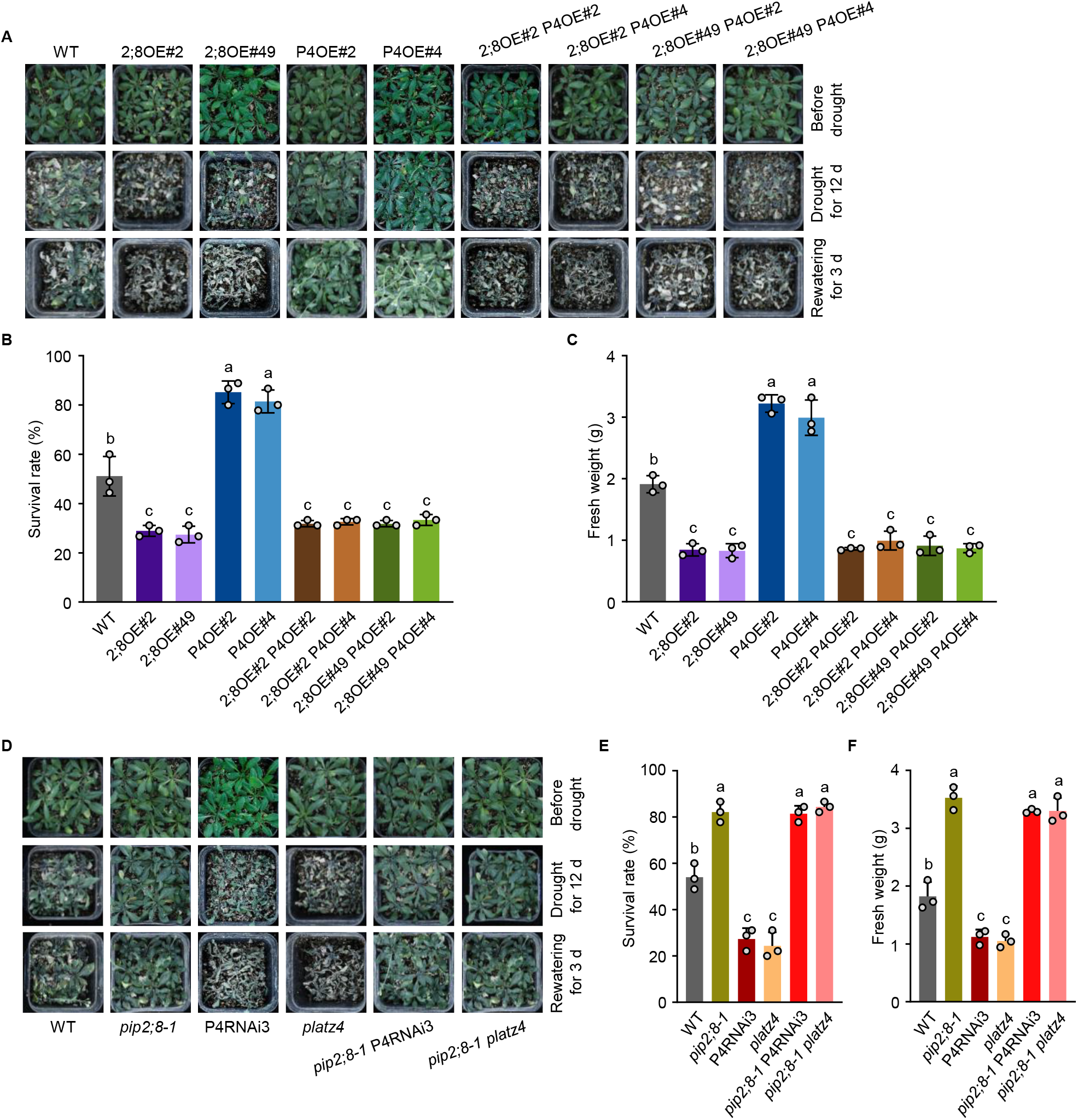
PIP2;8 functions epistaticaly to PLATZ4 in regulating drought tolerance in *Arabidopsis*. (A) Drought tolerance assays of WT, 2;8OE#2, 2;8OE#38, P4OE#2, P4OE#4, and 2;8OE#2 P4OE#2, 2;8OE#2 P4OE#4, 2;8OE#49 P4OE#2 and 2;8OE#49 P4OE#4 plants. Three-week-old plants were subjected to drought stress by no watering for 12 d and then rewatered for 3 d. (B,C) Survival rates and fresh weight of plants in (A), respectively, which were recorded 3 d after rehydration. Error bars indicate SEM (*n* = 3). Bars labeled with different lowercase letters are significantly different from one another (*P* < 0.05; one-way ANOVA). (D) Drought tolerance assays of WT, *pip2;8-1*, P4RNAi3, *platz4*, and *pip2;8-1* P4RNAi3, *pip2;8-1 platz4* plants. Three-week-old plants were subjected to drought stress by no watering for 12 d and then rewatered for 3 d. (E,F) Survival rates and fresh weight of plants in (D), respectively, which were recorded 3 d after rehydration. Error bars indicate SEM (*n* = 3). Bars labeled with different lowercase letters are significantly different from one another (*P* < 0.05; one-way ANOVA).

To obtain more information about how PLATZ4 and PIP2;8 are involved in the response to drought stress and ABA, we monitored the expression levels of *PLATZ4* and *PIP2;8* by RT-qPCR (Supplemental Figure 11). The results revealed that expression of *PLATZ4* was decreased by mannitol and ABA, whereas that of *PIP2;8* was induced by both treatments. These results support the idea that PLATZ4 inhibits the expression of *PIP2;8*.

### PIP2;8 shows water channel activity and involves in ABA response and stomatal closure regulation

Although PIPs in *Arabidopsis* have been shown to have aquaporin activity, PIP2;8 has not previously tested in this regard. Hence, we performed swelling experiments in *Xenopus* oocytes cells in hyper-osmotic solution. The swelling rate was lower in *Xenopus* oocytes injected with 46 ng cRNA of *PIP2;8* than in oocytes injected with water (negative control) (Figure 6A), indicating water efflux activity under this condition. S271 and S274 are two serine amino acids whose phosphorylation were changed by CEPR2 overexpression in Arabidopsis(Zhang et al., 2021). Two mutated cRNAs of *PIP2;8*, including *PIP2;8^S271AS274A^* and *PIP2;8^S271DS274D^* were also used in this experiment (Supplemental Figure 12). The swelling rate of *Xenopus* oocytes injected with 46 ng cRNA of *PIP2;8^S271AS274A^* was lower than that of cells injected with *PIP2;8* cRNA, but higher in cells injected with an equal amount of *PIP2;8^S271AS274D^* cRNA, indicating that phosphorylation at S271 or S274 activated the water channel activity of PIP2;8 (Figure 6A).

**Figure 6.**
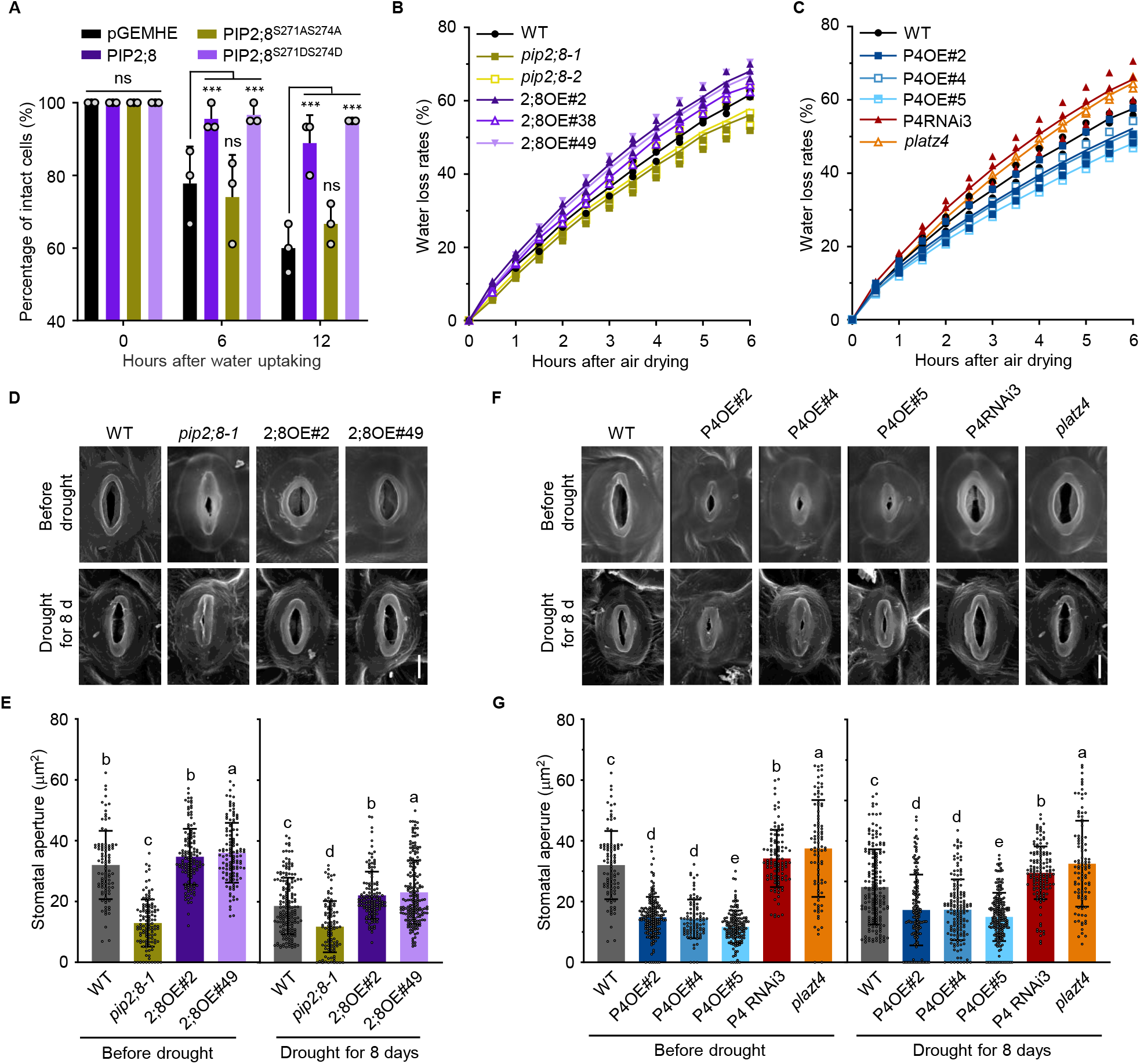
PIP2;8 shows water channel activity and involves in stomatal closure regulation. (A) Measurement of water channel activity in *Xenopus oocytes* cells expressing PIP2;8, PIP2;8^S271AS274A^ or PIP2;8^S271DS274D^. Xenopus oocytes injected with water were used as the negative control. S271AS274A and S271DS274D represent serine at sites of 271 and 274 were mutant to Alanine and Aspartate, respectively. Error bars indicate SEM (*n* = 3). ns, not significant, ***, *P* < 0.001 (one-way ANOVA). (B) Water loss rates of WT, *pip2;8-1, pip2;8-2* and 2;8OE2, 2;8OE38, 2;8OE49 plants. Aerial parts of 3-week-old plants were excised and weighed at different time points after detachment. Values are means ± SD of nine individual plants per genotype. Repeated experiments showed similar results. (C) Water loss rates of WT, P4OE#2, P4OE#4, P4OE#5, P4RNAi3, and *platz4* plants. Aerial parts of 3-week-old plants were excised and weighed at different time points after detachment. Values are means ± SD of nine individual plants per genotype. Repeated experiments showed similar results. (D) Representative photographs of stomatal of WT, *pip2;8-1*, 2;8OE#2, and 2;8OE#49 plants. Three-week-old plants were subjected to drought stress with or without watering for 8 d. Rosette leaves were used to observe the stomata with scanning electron microscope. bar = 5 μm. (E) Comparison of drought-induced stomatal closure in plants in (D). Values represent means ± SD (*n* > 50) from three biological replicates. Bars labeled with different lowercase letters are significantly different from one another (*P* < 0.05; one-way ANOVA). (F) Representative photographs of stomatal in WT, P4OE#2, P4OE#4, P4OE#5, P4RNAi3, and *platz4* plants. Three-week-old plants were subjected to drought stress with or without watering for 8 d. Rosette leaves were used to observe the stomata with scanning electron microscope. bar = 5 μm. (G) Comparison of drought-induced stomatal closure in plants in (F). Values represent means ± SD (*n* > 50) from three biological replicates. Bars labeled with different lowercase letters are significantly different from one another (*P* < 0.05; one-way ANOVA).

Next, we investigated the function of PIP2;8 in yeast cells. Under mannitol stress, the growth rate of PIP2;8-transformed cells were faster than control cells (yeast transformed with empty vector) (Supplemental Figure 13). The growth rate of PIP2;8^S271DS274D^-transformed yeast cells was similar to those of PIP2;8-transformed cells, whereas the growth rate of PIP2;8^S271AS274A^-transformed yeast cells was similar to the control. However, no transformed yeast cells exhibited a growth difference under normal condition (Supplemental Figure 13). These data indicate that phosphorylated PIP2;8 has water influx activity under mannitol stress.

We also measured the water loss of PIP2;8 and PLATZ4-overexpressing and mutant plants. The water loss of 2;8OE seedlings was higher than that of WT, whereas that of *pip2;8-1* and *pip2;8-2* mutants was lower (Figure 6B). Correspondingly, the stomatal aperture of 2;8OE plants was larger than WT, whereas those of *pip2;8-1* and *pip2;8-2* mutants were smaller (Figure 6D-6E). By contrast, water loss of P4OE seedlings was lower than WT, whereas that of P4RNAi3 and *platz4* mutants was higher (Figure 6C). Water loss and stomatal aperture of 2;8OE plants were similar to those of P4RNAi3 and*platz4* mutant, whereas *pip2;8-1* and*pip2;8-2* mutants were similar to P4OE plants (Figure 6B-6G). These data illustrate that PIP2;8 functions as an effective water channel to negatively regulate stomatal closure, which is essential for the function of PLATZ4.

### AITR6 interacts with PLATZ4 to promote the inhibitory effect of PLATZ4 on *PIP2;8* expression

To explore the regulation of PLATZ4 in response to ABA and drought stress, we examined the interaction of PLATZ4 and AITR6, which interacts with PLATZ4 in yeast two-hybrid (Y2H) assays(Liu *et al*., 2020). In the Y2H test, yeast cells co-transformed with the PLATZ4-BD and AITR6-AD constructs were able to grow on synthetic medium lacking Trp, Leu, His, and 25 mM AbA (aureobasidin A) (quadruple dropout [QDO]), confirming the physical interaction of PLATZ4 with AITR6 (Figure 7A). We further confirmed the interaction between PLATZ4 and AITR6 by BiFC and pulldown assays (Figure 7B and 7C).

**Figure 7.**
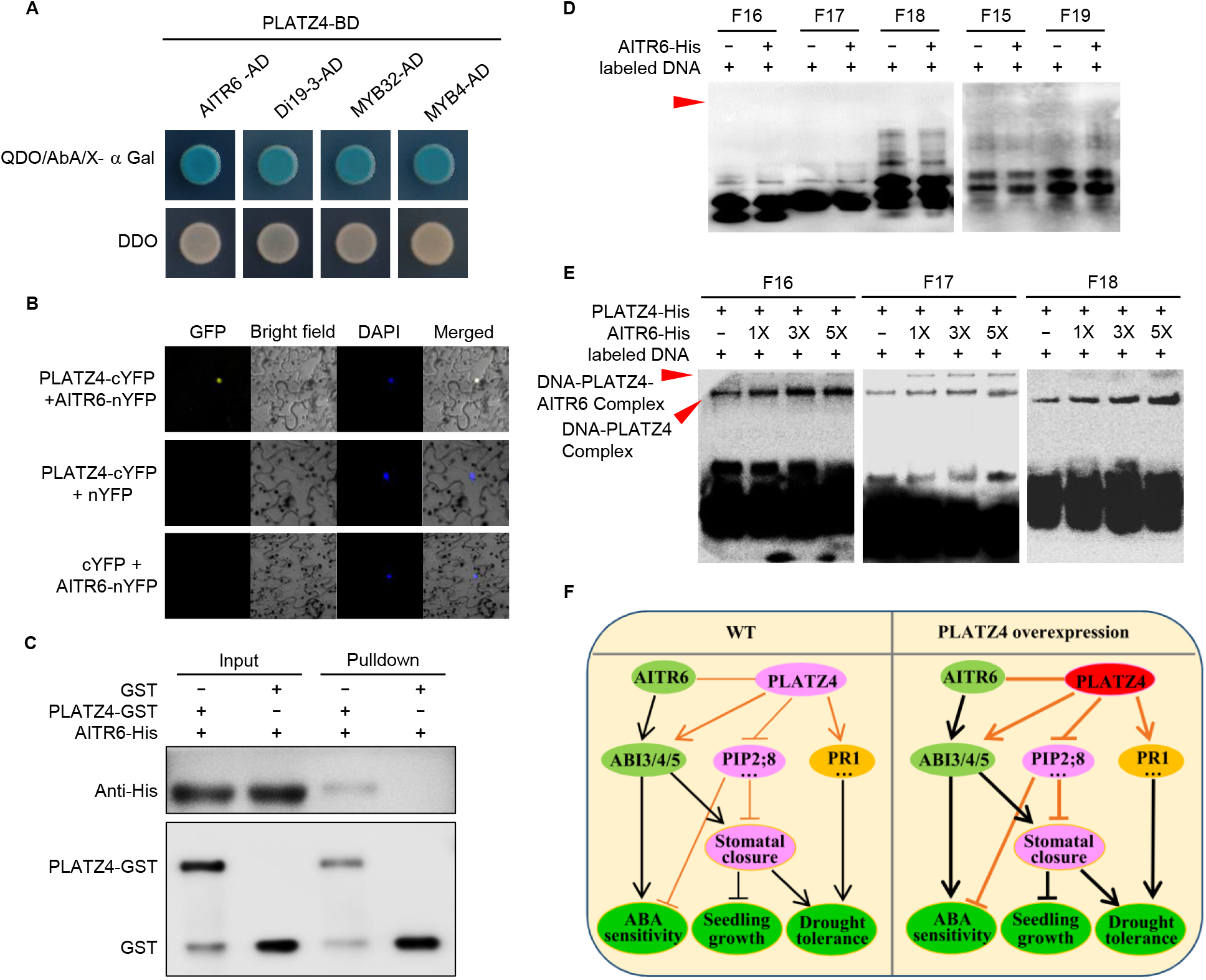
AITR6 interacts with PLATZ4 to promote the inhibitory effect of PLATZ4 on *PIP2;8* expression. (A) Yeast two-hybrid assays showing the interaction between PLATZ4 and AITR6, Di19-3, MYB32, and MYB4. DDO, Synthetic Dropout/-Leu-Trp; QDO, Synthetic Dropout/-Leu-Trp-His-Ade. (B) Bimolecular fluorescence complementation (BiFC) assays showing the interaction between PLATZ4 and AITR6 in *N. benthamiana*. DAPI, Nuclear dye, 4′,6-diamidino-2-phenylindole. (C) In vitro pulldown assays showing the interaction between PLATZ4 and AITR6. AITR6-His was incubated with GST or GST-PLATZ4 purified from *E. coli* and was shed to remove unbound proteins. The bound proteins were eluted and analyzed by immunoblotting with anti-GST and anti-His antibodies. (D) EMSA analysis of the binding activity of AITR6 to the F15-F19 DNA fragments in the *PIP2;8* promoter. The plus (+) and minus (−) signs denote the presence or absence of the protein or DNA fragments in each sample. (E) EMSA analysis of the binding activity of PLATZ4 to the F16, F17 and F18 DNA fragments in the PIP2;8 promoter in the presence or absence of AITR6. The plus (+) and minus (−) signs denote the presence or absence of the protein or DNA fragments in each sample. (F) Work models for AtPLATZ4 in positively regulating ABA sensitivity and plant drought tolerance but negatively regulating plant growth. In which, PLATZ4 unpregulates the expression of *ABI3, ABI4* and *ABI5*, key genes involved in ABA signaling, which results in ABA sensitivity and stomatal closure. Moreover, PLATZ4 inhibits the expression of *PIP2;8* and other stress related genes to reduce water movement, which results in stomatal closure. Under normal condition, stomatal closure results plant growth inhibition; Under drought stress, stomatal closure increases plant drought tolerance. Besides, PLATZ4 unpregulates the expression of *PR1* to increase drought tolerance. PLATZ4 fulfils these function by interacting with AITR6, which inreases the binding of *PIP2;8* promoter by PLATZ4.

Next, we analyzed the binding of AITR6 to the *PIP2;8* promoter. The results revealed that AITR6 did not directly bind the three DNA fragments within the *PIP2;8* promoter mentioned above (Supplemental Figure 14). EMSA assays revealed no binding of AITR6 to the three probes (F16, F17, and F18) bound by PLATZ4 or the two probes (F15 and F19) that PLATZ4 did not bind (Figure 7D). However, higher concentrations of AITR6 enhanced the binding of PLATZ4 to the F16, F17 and F18 DNA fragments (Figure 7E). These data suggest that AITR6 acts as a corepressor of PLATZ4 in the inhibition of *PIP2;8* expression.

## Discussion

Drought is a major abiotic stress affecting agricultural production around the world. Accordingly, research on the development of drought-tolerant crop plants is of critical importance. Accumulating evidence has revealed the important roles of PLATZs in many processes of plant growth, development, and responses to environmental stimuli, including regulation of the stem cell niche and root meristem size(Yamada *et al*., 2020), endosperm development and seed filling(Li *et al*., 2017; Wang *et al*., 2019a), leaf cell proliferation and leaf senescence(Kim *et al*., 2018), and desiccation tolerance in seeds(González-Morales *et al*., 2016). PLATZ1 participates in drought tolerance in vegetative tissues(González-Morales *et al*., 2016), and PLATZ2 negatively regulates salt tolerance(Liu *et al*., 2020; Zhang *et al*., 2018). *PLATZ4* was significantly inhibited by mannitol and ABA in *Arabidopsis* (Supplemental Figure 11). Seedlings overexpressing *PLATZ4* improved drought tolerance, while RNAi of *PLATZ4* or *platz4* mutation decreased it (Figure 1), indicating that *PLATZ4* positively regulates drought tolerance. Therefore, we provide a novel transcriptional regulator of plant drought tolerance.

Aquaporins (AQP) are primarily involved in maintaining cellular water homeostasis(Wang *et al*., 2020; Zargar *et al*., 2017). Several lines of evidence in various crop plants indicate that the AQPs are responsible for precise regulation of water movement and consequently play a crucial role in the drought tolerance(Bienert and Bienert, 2017). We found that the expression of *PIP2;8*, which encodes a plasma membrane AQP, was significantly down-regulated by *PLATZ4* overexpression but up-regulated by *PLATZ4* suppression (Figure 3A and Supplemental Figure 7A), revealing a role of PLATZ4 in suppressing the expression of *PIP2;8*. Observations of GUS activity driven by *PIP2;8* promoter in the presence of PLATZ4 and GFP (Supplemental Figure 8A), as well as Y1H (Figure 3B), Dual-Luciferase Reporter Assay (Figure 3C), ChIP-qPCR (Figure 3D), and EMSA assays (Figure 3E), confirmed the binding of PLATZ4 to the A/T-rich sequences between −827 bp and −728 bp within *PIP2;8* promoter. The binding to A/T-rich sequence and repressive features of PLATZ4 are consistent with previous studies reporting non-specific A/T-rich sequence binding and repressive activity of PLATZs against their target genes(Liu *et al*., 2020; Nagano *et al*., 2001). Importantly, the responses of *PIP2;8*-overexpressing plants and the *pip2;8* mutant to drought stress and ABA were opposite to that of PLATZ4-overexpressing plants and the *platz4* mutant, respectively (Figure 3), confirming that PLATZ4 inhibits *PIP2;8*. Further genetic analysis demonstrated that PLATZ4 acts as a novel transcription repressor of *PIP2;8* to positively regulate drought tolerance in *Arabidopsis* (Figure 5). Although numerous environmental factors such as drought, salinity, cold, and light regulate AQPs at the transcriptional level(Deshmukh *et al*., 2017; Liu *et al*., 2013a), only a few transcription factors, including ABI3(Mao and Sun, 2015), ASR1(Ricardi *et al*., 2014), and MdDREB6.2(Wang *et al*., 2017), have been reported to directly bind the promoters of *AQP* genes. Hence, we uncovered a novel transcriptional regulator of *AQP* genes in response to drought stress.

In addition to *PIP2;8*, other genes were transcriptionally regulated by PLATZ4, including At1g33840, At3g01345, *CYSTM8*, and *PR1*(Supplemental Figure 7 and Supplementary Table 1). *PR1* was up-regulated by 20- and 100-fold by *PLATZ4* overexpression under normal and drought stress, respectively. And *PR1*-overexpressing plants exhibit drought tolerance(Liu et al., 2013b). *Arabidopsis* Di19 (Drought-induced, At1g56280), a transactivator, directly binds to the promoters of *PR1, PR2*, and *PR5* to increase their expressions(Liu *et al*., 2013b). Di19-3, as well as AITR6, MYB32, MYB3, MYB7, NAC066, ARIA, and AFP3, physically interact with all of PLATZ family proteins in the Y2H assay(Liu *et al*., 2020; Trigg et al., 2017) (Figure 7A). These results indicated that PLATZ4 might regulate *PR1* expression by interacting with Di19. Overexpression of *OsDi19-4, TaDi19A, GhDi19-2*, and *GmDi19-5* causes hyper-sensitivity to ABA and tolerance to drought stress(Feng et al., 2015; Li et al., 2010a; Li et al., 2010b; Wang et al., 2016). Here, we confirmed that AITR6 interacts with PLATZ4 by Y2H, BiFC, and pulldown assays (Figure 7A-7C). Although AITR6 did not bind the *PIP2;8* promoter directly (Figure 7D and Supplemental Figure 12), it did enhance the *PIP2;8* promoter binding by PLATZ4 (Figure 7D), indicating that AITR6 acts as a transcriptional co-suppressor of PLATZ4 in the regulation of *PIP2;8* expression. Taken together, these findings indicate that PLATZ4 enhances drought tolerance by interacting with other transcriptional regulators to suppress or increase the expression of target genes.

The ABA-induced transcription repressor AITR6 is involved in ABA and drought response in *Arabidopsis(Song et al*., 2016; Tian *et al*., 2017), indicating a possible role for *PLATZ4* in the ABA response in *Arabidopsis*. In fact, *PLATZ4-*overexpressing plants were hyper-sensitive, whereas RNAi and loss-of-function plants were insensitive to ABA during both germination and later growth stages (Figure 7 and Supplemental Figure 2). Although PLATZ4 did not bind the *ABI3, ABI4*, or *ABI5* promoters (Supplemental Figure 4), it up-regulated the expression of all three genes (Supplemental Figure 3). This indicated a positive role of PLATZ4 in ABA signaling, which is directly connected to the plant water status and stomatal closure(Joshi-Saha et al., 2011). ABA-triggered stomatal closure requires an increase in guard cell permeability to water and possibly hydrogen peroxide through OST1-dependent phosphorylation of PIP2;1 at Ser-121(Grondin et al., 2015). Aquaporin function in stomata movement is specifically associated with water stress (hyperosmotic treatment)(Chen et al., 2021; Shope and Mott, 2006). Here, we demonstrated the water channel activity of PIP2;8 by swelling assays of *Xenopus* oocytes (Figure 6A), the increased tolerance of PIP2;8-transformed yeast cells to mannitol stress (Supplemental Figure 13), and the tolerance of the *pip2;8* mutant to drought stress (Figure 4). The increased stomatal closure of *pip2;8* and the increased stomatal opening of PIP2;8 overexpression plants relative to WT were observed under normal and drought stress (Figure 6D-6E). These data indicate a role for PLATZ4-PIP2;8 module in regulation of water status-mediated stomatal closure.

Based on these findings, we propose the model shown in Figure 7 F. Under normal conditions, when PLATZ4 is overexpressed, *ABI3, ABI4*, and *ABI5* are upregulated, whereas *PIP2;8, DEG3*, and At3g01345 are downregulated relative to WT. These effects lead to ABA sensitivity and stomatal closure, which in turn results in inhibition of seedling growth. Under drought stress, when PLATZ4 is overexpressed, *ABI3, ABI4*, and *ABI5* are further up-regulated, whereas *PIP2;8*, DEG3, and At3g01345 are further inhibited, leading to improved ABA sensitivity, further stomatal closure, and then drought tolerance. Moreover, PLATZ4 interacts with AITR6 to inhibit the expression of *PIP2;8* to promote stomatal closure, resulting in drought tolerance, and activate the expression of *ABI3, ABI4*, and *ABI5* to increase ABA sensitivity. In addition, PLATZ4 upregulates *PR1* to increase drought tolerance. Therefore, promoting the expression of *PLATZ4* or inhibiting the expression of *PIP2;8* represents a new strategy for improving plant drought tolerance.

## Materials and Methods

### Plant materials and growth conditions

*Arabidopsis thaliana* (L.) Heynh. cv. ‘Columbia-0’ was used as the WT. *Arabidopsis* plants were grown as previously described(Chen et al., 2014). The T-DNA insertion lines Salk_099098 (*platz4*), Salk_099098 (*pip2;8-1*), and SK16840 (*pip2;8-2*), all in the Col-0 background, were obtained from the *Arabidopsis* Biological Resource Center (http://www.arabidopsis.org). Homozygous plants were verified by RT-PCR with primers listed in Supplementary Table 2. PLATZ4 and PIP2;8–overexpressing lines were generated by introducing their coding regions into *pBI121/pSuper1300* vector under the control of the cauliflower mosaic virus 35S promoter. To generate a *PLATZ4* RNAi construct, the pFGC5941 vector for dsRNA production was used. The transformation of *Arabidopsis* was performed by floral dip using *Agrobacterium tumefaciens* strain GV3101. T_3_ transgenic plants were selected on half-strength MS medium (1.5% sucrose and 0.85% agar) supplemented with 50 mg/L kanamycin or 25 mg/L homomycin B and confirmed by RT-PCR using primers listed in Supplementary Table 2. The 2;8OE#2 P4OE#2, 2;8OE#2 P4OE#4, 2;8OE#49 P4OE#2, 2;8OE#49 P4OE#4, *pip2;8-1* P4RNAi3, *pip2;8-1 platz4* were generated by crossing, and F_3_ homozygous lines were genotyped and kept for further study. All selected *Arabidopsis* plants were grown in a greenhouse under a 16 h light/8 h dark cycle at 22°C with a light intensity of 9600 LUX.

### Phenotype analysis

Plants of different genotypes were grown under the same conditions in a greenhouse. Seeds were collected at the same time. To observe seedling’s phenotypes, plants were grown for approximately 2 weeks under normal watering conditions and subjected to drought stress (no watering) for the indicated days. Drought phenotype analysis was performed as described by liu et al.(Liu *et al*., 2013b). In each experiment, nine plants were grown in a small pot under a 16 h light/ 8 h dark photoperiod at 22°C; at least three independent experiments were performed. Photos were taken after 8 or 12 d of drought treatment. The plants were re-watered and photographed 3 d later, and survival rates were calculated.

For the water loss assay, rosette leaves of 3-week-old plants were detached, placed on filter paper, and left on the bench at room temperature. Fresh weight was monitored at the indicated times. Each phenotype was performed with four replicate samples, and each sample contains 10 mature leaves from 10 different plants.

For ABA response during the seed germination stage, 1 or 2 μM ABA was added to half-strength MS medium for 8 or 10 d. For ABA response after germination, 36 3-day-old seedlings for each genotype were transferred onto half-strength MS medium containing 5 or 10 μM ABA for 14 d. The root lengths of the treated seedlings were measured using a ruler. The fresh weights of whole seedlings were measured using a four-decimal place balance. All experiments were performed with six biological replicates.

PIP2;8, PIP2;8^S271DS274D^, PIP2;8^S271AS274A^ were cloned into pYES2, and the resultant constructs were transformed into *Saccharomyces cerevisiae* strain YPH500. Yeast transformants were spotted on the selective medium without Ura, and positive clones were cultured on medium without Ura containing galactose (20%), raffinose (20%), and BU-salt (50 ml: 1.95 g Na_2_HPO_4_, 1.855 g NaH_2_PO_4_·2H_2_O). Positive clones were cultured in liquid medium with or without 2 M mannitol at 30°C for 36 h. Optical densities (OD) at 600 nm were recorded for independent transformants. Error bars indicate SEM (n = 3). *, *P* < 0.05; **, *P* < 0.01; ***, *P* < 0.001 (one-way ANOVA).

#### RNA extraction, RT-PCR, and quantitative RT-PCR (RT-qPCR)

Plants used to detect the expression levels of related genes were grown on half-strength MS medium for 14 d, and then transferred to half-strength MS liquid medium containing 300 mM mannitol or 30% PEG6000 or 50 μM ABA or not for indicated times. Samples were collected at the indicated time points in liquid nitrogen and stored at −80°C.

Total RNA from the above samples was extracted using RNAiso Plus (Takara, Ohtsu, Japan). As described by Yu *et al*(Yu et al., 2019), reverse transcriptions were performed using PrimeScriptTM RT reagent kit (TaKaRa, Ohtsu, Japan). The RT-qPCR analyses were performed using the TransStart Tip Green qPCR SuperMix (TransGen, Beijing, China) and CFX96 (Bio-Rad, Hercules, CA, USA). *GAPDH* and *UBQ10* were used as the internal controls. The cDNA used for RT-PCR was synthesized using the PrimeScrip First-Strand cDNA Synthesis Kit (TaKaRa). RT-PCR cycling conditions were as follows: denaturation at 94°C for 5 min, followed by 23–34 cycles of amplification, and final elongation at 72°C for 5 min. *EF-1α* was used as the internal control. Primers are listed in Supplementary Table 2.

### Fluorometric GUS assays

The fluorometric GUS assays were performed as previously described(Li et al., 2010c). *pPIP2;8* representing the 1813 bp intergenetic region upstream of ATG of *PIP2;8*, and was fused to the *pBI121* vector to drive *GUS* expression. GUS enzymatic activity resulted from transiently transformed *N. benthamiana* leaves using the constructs *ProPIP2;8:GUS* and *Pro35S:PLATZ4, Pro35S:AITR6*, and *Pro35S:GFP* (negative control) was measured on an F-4500 fluorescence spectrofluorometer (Hitachi, Tokyo, Japan) using 4-methylumbelliferyl-β-d-glucuronide as a substrate. Standard curves were prepared with 4-methylumbelliferone. Average GUS activity was obtained from three independent transformants, and each assay was repeated three times.

### Y2H and Y1H assays

The matchmaker GAL4 two-hybrid system (Clontech) was used for Y2H assays. Full-length *PLATZ4* and *AITR6* were cloned into the pGBKT7 (BD) and pGADT7 (AD) vectors, respectively. Primers used are listed in Supplementary Table 2. Constructs were co-transformed into the yeast strain AH109. Medium supplemented without Leu or Trp (double dropout [DDO]) was used for growth and Leu-Trp-His-Ade (QDO) for selection.

The F1, F2, and F3 promoter regions of *PIP2;8* were amplified and cloned into pLacZi2μ, which contains the LacZ reporter gene, to generate *ProF1:LacZ, ProF2:LacZ*, and *ProF3:LacZ*, respectively. The promoter regions of *ABI3, ABI4, ABI5*, and *PIP2;6* were also amplified and cloned into the pLacZi2μ vector. The CDS of *PLATZ4* and *AITR6* were cloned into the pJG4-5 vector containing the GAL4 transcriptional activation domain (GAD) to generate PLATZ4-pJG4-5 and AITR6-pJG4-5. The two types of plasmids were co-transformed into yeast strain EGY48 according to the Yeast Protocols Handbook (Clontech) as described previously(Lin et al., 2007). Cells were plated onto the first selective medium without Ura and Trp, and positive clones were cultured on medium without Ura and Trp containing galactose (20%), raffinose (20%), BU-salt (50 ml: 1.95 g Na_2_HPO_4_, 1.855 g NaH_2_PO_4_·2H_2_O), and 5-bromo-4-chloro-3-indolyl -β-D-galactopyranoside (X-Gal) for development of blue color(Zhuang et al., 2019). Cells co-transformed with pJG4-5 and *ProF1:LacZ, ProF2:LacZ*, or *ProF3:LacZ* were used as negative controls.

### BiFC Assays

The coding regions of AITR6 and PLATZ4 were amplified and cloned into pSPYNE-35S and pSPYCE-35S, which contain DNA encoding the N- or C-terminal fusions of YFP (nYFP or cYFP), respectively. The two resulting constructs and the cotransformation vector *Pro35S:P19* were transformed into the *Agrobacterium tumefaciens* GV3101 strain. The *A. tumefaciens* strains were incubated, harvested, and resuspended in infiltration buffer (10 mM MES, 0.2 mM acetosyringone, and 10 mM MgCl_2_) at a final concentration of OD_600_ = 1.5. Equal volumes of different combinations of *Agrobacterium* strains were mixed and injected into *N. benthamiana* leaves using a needleless syringe. The leaves were cultivated at 24 °C for 72 h prior to the detection of YFP fluorescence. Primers used in this experiment are listed in Supplemental Table 2.

### Pulldown assays

As described previously(Yu *et al*., 2019), characterizing the interaction between PLATZ4 and AITR6 *in vitro*, the CDS of *PLATZ4* was fused with GST in vector pGEX-4T-3 to yield *PLATZ4-GST*, and the full-length CDS of *AITR6* was fused with the His tag in pET30a vector to generate *AITR6-His*. The resultant constructs were transformed into competent *E. coli* Rosetta cells. Protein expression was induced with 1 mM isopropyl β-D-thiogalactoside (IPTG) for 12 h at 16°C. The AITR6-His protein was purified using the His-Tagged Protein Purification Kit (CWBIO, Beijing, China) and PLATZ4-GST was purified using BeyoGold GST-tag Purification Resin (BeyoGold, Shanghai, China). In the pulldown assay, 25 μg of PLATZ4-GST, 25 μg of AITR6-His, and BeyoGold GST-tag Purification Resin were incubated in 1 ml of GST pull-down binding buffer (50 mM Tris-HCl 200 mM NaCl 1mM EDTA 1% [v/v] NP-40 1 mM DTT, and 10 mM MgCl_2_ [pH 8.0]) at 4°C for 2 h. After incubation, GST proteins were eluted with GST-pulldown binding buffer (50 mM Tris-HCl 400 mM NaCl 1 mM EDTA 1 mM DTT and 50 mM GSH [pH 8.0]) and analyzed using anti-His antibodies (TransGen Biotech, Beijing, China). Primers used in this experiment are listed in Supplemental Table 2.

### Dual-Luciferase Reporter Assays

Dual-Luciferase Reporter Assays were performed as described previously(Liu et al., 2019). To construct the *ProPIP2;8:LUC* plasmids, we amplified the F3–F9 promoter regions of *PIP2;8* (Figure 3B and Supplemental Figure 8B) and the 2;6F1–2;6F3 promoter regions of *PIP2;6* (Supplemental Figure 9B) and cloned into the pGreenII0800-LUC to generate reporters. The full-length cDNA of *PLATZ4* was cloned into vector pBI121 to construct *Pro35S:PLATZ4*, which was used as the effector. *Pro35S:GFP* was used as the negative control. LUC and REN activities were measured by the Dual-Luciferase Reporter Assay System (Promega Corporation, Madison, WI, USA), and LUC activity was normalized against REN activity. Primers used for these constructs are listed in Supplemental Table 2.

### Chromatin immunoprecipitation (ChIP)-qPCR assays

The ChIP assays were performed as described previously(Du et al., 2019). Seven-day-old *ProSuper:PLATZ4-HA* transgenic seedlings and HA-specific monoclonal antibody (Beyotime Biotechnology, China) were used for ChIP. The ChIP DNA products were analyzed by qPCR with primers designed to amplify the F10–F14 DNA fragments of the of *PIP2;8* promoter (Supplemental Figure 8C) and 2;6F4–2;6F6 DNA fragments in the promoters of *PIP2;6*, respectively (Supplemental Figure 9B).

### EMSA assays

In this experiment, the PLATZ4-GST and AITR6-His proteins were induced in *E.coli* Rosetta cells and purified as described by Yu et al.(Yu *et al*., 2019). The AITR6-His protein was purified using the His-Tagged Protein Purification Kit (CWBIO, Beijing, China) and PLATZ4-GST was purified using BeyoGold™ GST-tag Purification Resin (BeyoGold, Shanghai, China). Five DNA sequences (F14-F19) within the promoter of *PIP2;8* (Supplemental Figure 8D) were amplified using biotin-labeled primers (Supplementary Table 2) synthesized by Sangon Biotech (Shanghai, China) and purified by a PCR purification kit (Qiagen). The EMSA was conducted as described previously(An et al., 2018) using a LightShift Chemilumiescent EMSA Kit (Thermo Fisher Scientific, Pierce, USA). Unlabeled DNA fragments of the same sequences were used as competitors. Migration of biotin-labeled probes was detected using the enhanced chemiluminescence substrate (Thermo Scientific) on a ChemDoc XRS system (Bio-Rad).

### Fv/Fm detemination

The maximum photochemical efficiency of PSII [Fv/Fm, the ratio of variable (Fv) to maximum (Fm) fluorescence; (Fv/Fm = (Fm - F_0_)/Fm)] in the drought and control groups of WT, mutants, and OE lines were determined using a Handy PEA fluorimeter (FMS-2, Hansatech, UK)(Krause, 1991). The leaves were dark-adapted for 20 min before measurement. Sixteen leaves were tested for each treatment.

### Water channel activity

For the *Xenopus* oocytes uptake assay, the CDS of *PIP2;8, PIP2;8^S271AS274A^* and *PIP2;8^S271DS274D^* was amplified from *Arabidopsis* cDNA and cloned into destination vector pGEMHE. pGEMHE containing *PIP2;8, PIP2;8^S271AS274A^* or *PIP2;8^S271DS274D^* was linearized using *Sal*I (Takara, Beijing, China), and the cRNA was transcribed in vitro using the mMESSAGE mMACHINE T7 mRNA synthesis kit (Ambion). The cRNA quality was checked by agarose gel electrophoresis. *Xenopus oocytes* were isolated in 25 mL of ND96 free Ca^2+^ solution containing 43 mg of collagenase and 12.5 mg of trypsin inhibitor for 1.5 h. After isolation, cells were recovered in 25 mL of ND96 for another 24 h in 18°C. Every 15 *Xenopus* oocytes were then injected with 46 ng cRNA of empty vector, *PIP2;8, PIP2;8^S271AS274A^*, or *pip2;8^S271DS274D^*. Injected oocytes cultivated in ND96 with Ca^2+^ for 30 h were incubated in ddH_2_O for the indicated times. Intact cells were counted before and after the indicated times.

### RNA-Seq and data analysis

Three biological replicates of 2-week-old P4OE#2 and WT plants were grown on half-strength MS medium under long-day conditions (16 h light/8 h dark). Whole seedlings were treated with 30% PEG6000 as dehydration treatment or in a Petri dish with moistened Kim wipes as a control. The Petri dish was sealed with Parafilm and left for 4 h. Ten milligram of seedlings was considered to be one biological replicate. Total RNA was extracted using Trizol and RNeasy Mini Kit (Qiagen) with on-column DNase digestion. Library preparation and RNA-seq were performed using Capital Bio technology (Beijing, China) with an Illumina HiSeq 2000 with 50-bp single-end reads and 30 million reads per sample. Raw RNA-seq reads were subjected to quality checking and trimming. The trimmed reads of each sample were aligned to the publicly available reference genome and TAIR10 using GSNAP. The alignment coordinates of reads uniquely aligned to the reference genome were used for lookup, and read count tallies were computed for each annotated gene. Finally, RNA-seq reads were used to identify differentially expressed genes for comparison between P4OE#2 and WT plants subjected for control or dehydration treatment; this analysis was performed with the R package DESeq2. Normalization was conducted by DESeq2, which automatically corrects for biases introduced by differences in the total numbers of uniquely mapped reads in each sample. Normalized read counts were used to calculate fold changes and statistical significance (Data2Bio). Clustering was performed using the ‘aheatmap’ function of the NMF package in R and log2 reads per million mapped read values. Gene Ontology analysis was performed using the BiNGO software. Different expression genes between PLATZ4 overexpression and WT seedlings identified by RNA-Seq were shown in Supplemental Data 1.

### Accession numbers

Gene sequence data form this article can be found in the TAIR under the following accession number: PLATZ4 (At1g32700); PIP2;8 (At2g16850); EF-1α (At1g18070); UBQ10 (At4g05320); GAPDH (At1g16300); ABA1 (At5g67030); ABA2 (At1g52340); ABA3 (At1g16540); NCED3 (At3g14440); NCED9 (At1g78390); ABI1 (At4g26080); ABI3 (At3g26450); ABI4 (At2g40220); ABI5 (At2g36270); CYP707A2 (At2g29090); GUN4 (At3g59400); MYB41 (At4g28110); DEG3 (At4g29200); CEP5 (At5g66815); FRO5 (At5g23990); PRX2 (At1g05250); NRT2.3 (At5g60780); CYSTM8 (At3g22235); SIF1 (At1g51830); E12A11 (At1g18100); PR1 (At2g14610); PDF1.2 (At5g44420); PCR1 (At1g14880); SAG13 (At2g29350); PIP2;6 (At2g39010); MYB32 (At4g34990); MYB4 (At4g38620); AITR6 (At5g63350); Di19 (At1g56280); At1g19960; At1g33840; At1g68480; At3g01345; At3g60270; At5g42530; At5g46710. RNA-Seq raw data discussed in this study can be accessed at NCBI via BioProject ID: PRJNA764322.

## Supplemental Figures

**Supplemental Table 1** Different expression genes between *PLATZ4* overexpression and WT seedlings identified by RNA-Seq.

**Supplemental Table 2** Primers used in this study.

## Author contributions

C.W., C.Z., and S.Z. conceived the original screening and research plans; M.L., C.W., Z.J., and L.Z. performed experiments, analyzed the data, made the figures, and wrote several parts of the original article; C.L., J.H., G.Y., and K.Y. provided suggestions; C.W., C.Z., and S.Z. supervised and complemented the writing. All authors approved the final manuscript.

## Acknowledgments

This work was supported by the Natural Science Foundation of China (Grant number 31970292 and 31972357), the National Key R&D Program of China (2018YFD1000704/2018YFD1000700), and the Major Program of Shandong Province Natural Science Foundation (ZR2018ZB0212). We also thank Gristopher K Patil (San Francisco) for the linguistic assistance during the preparation of this manuscript.

## Conflict of interest

The authors declare that they have no conflict of interest.

## Data availability

All relevant data, vectors, and plant materials that support the findings of this study are available from the corresponding author upon request.

---

**Supplementary Figure 1.**
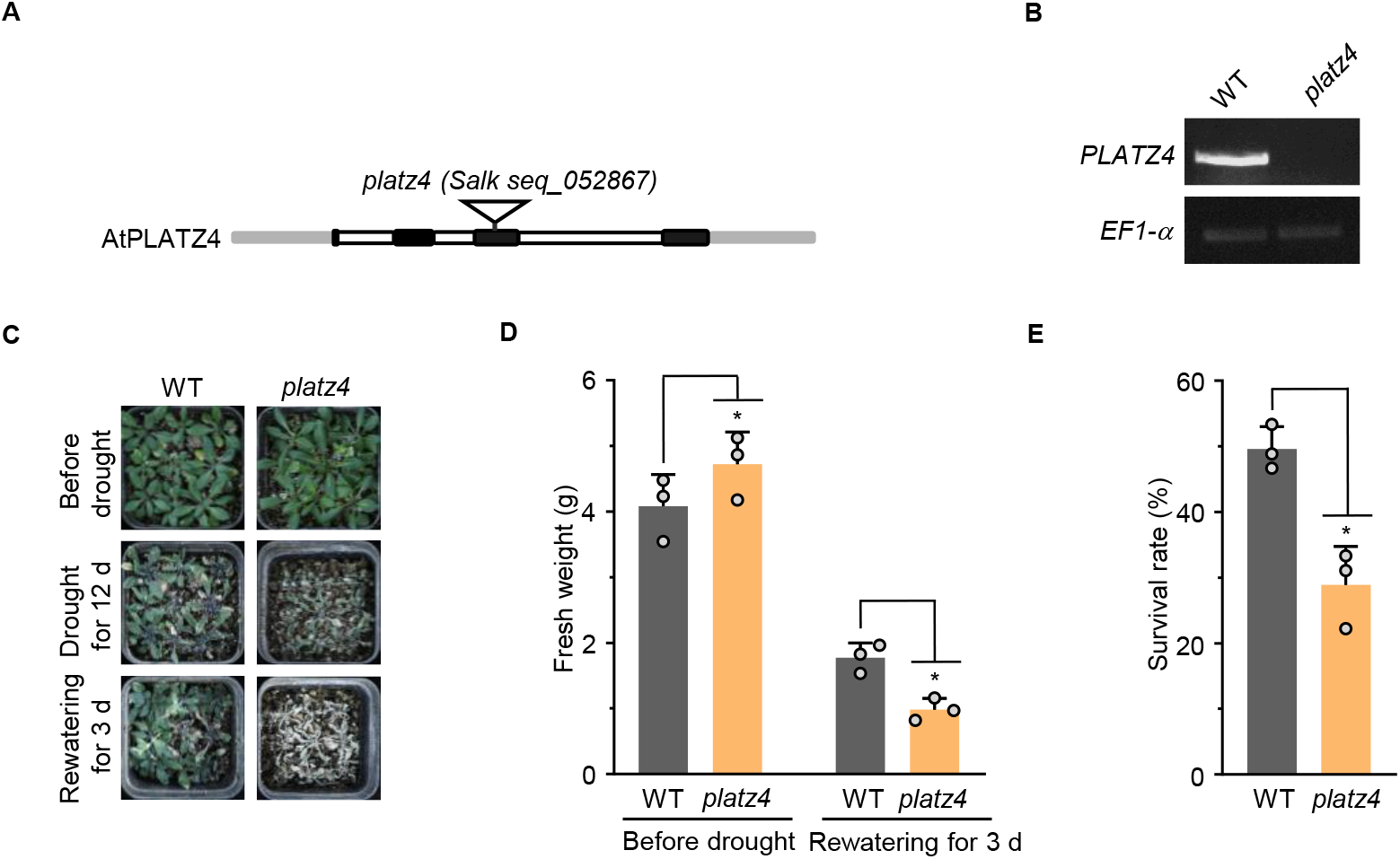
Drought tolerance assays of WT and *platz4* seedlings. (A) Schematic illustration of the T-DNA insertion site in the *platz4* mutant. (B) *PLATZ4* transcript levels in WT and *platz4* mutant as determined by RT–PCR. *EF-1a* was used as the internal. (C) Drought tolerance assay of WT and *platz4* seedlings. Three-week-old plants were subjected to drought stress by no watering for 12 d and then rewatering for 3 d. (D,E) Fresh weight (D) and survival rates of plants in (C). Fresh weight were recorded before drought and 3 d after rewatering, respectively. Survival rates were recorded 3 d after rewatering. Error bars indicate SEM (*n* = 3). *, *P* < 0.05 (Student’s t-test).

**Supplementary Figure 2.**
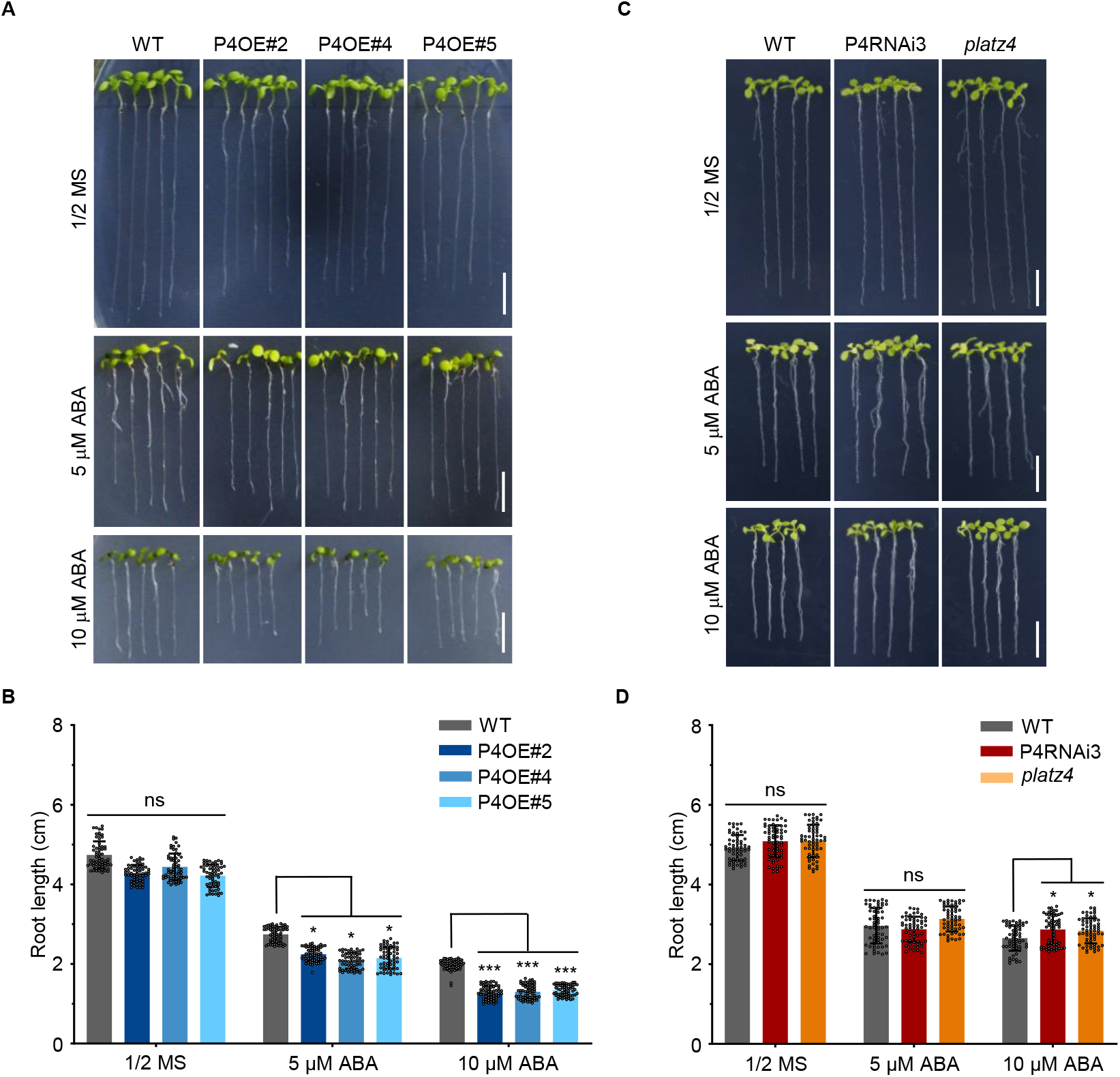
PLATZ4 positively regulates ABA response in Arabidopsis seedlings. (A) Phenotypes of 4-day-old uniformly developed seedlings of WT, P4OE#2, P4OE#4 and P4OE#5 plants grown on half–strength MS medium with or without 5 or 10 μM ABA for 10 d. (B) Length of the primary roots of the seedlings shown in (A). Error bars indicate SEM (*n* = 60). The asterisks indicate significant differences compared with WT. ns, not significant; *, *P* < 0.05; ***, *P* < 0.001 (one-way ANOVA). (C) Phenotypes of 4-day-old uniformly developed seedlings of WT, P4RNAi3 and *platz4* plants grown on on half–strength MS medium with or without 5 or 10 μM ABA for 10 d. (D) Length of the primary roots of the seedlings shown in (C). Error bars indicate SEM (*n* = 60). The asterisks indicate significant differences compared with WT. ns, not significant; *, *P* < 0.05 (one-way ANOVA).

**Supplementary Figure 3.**
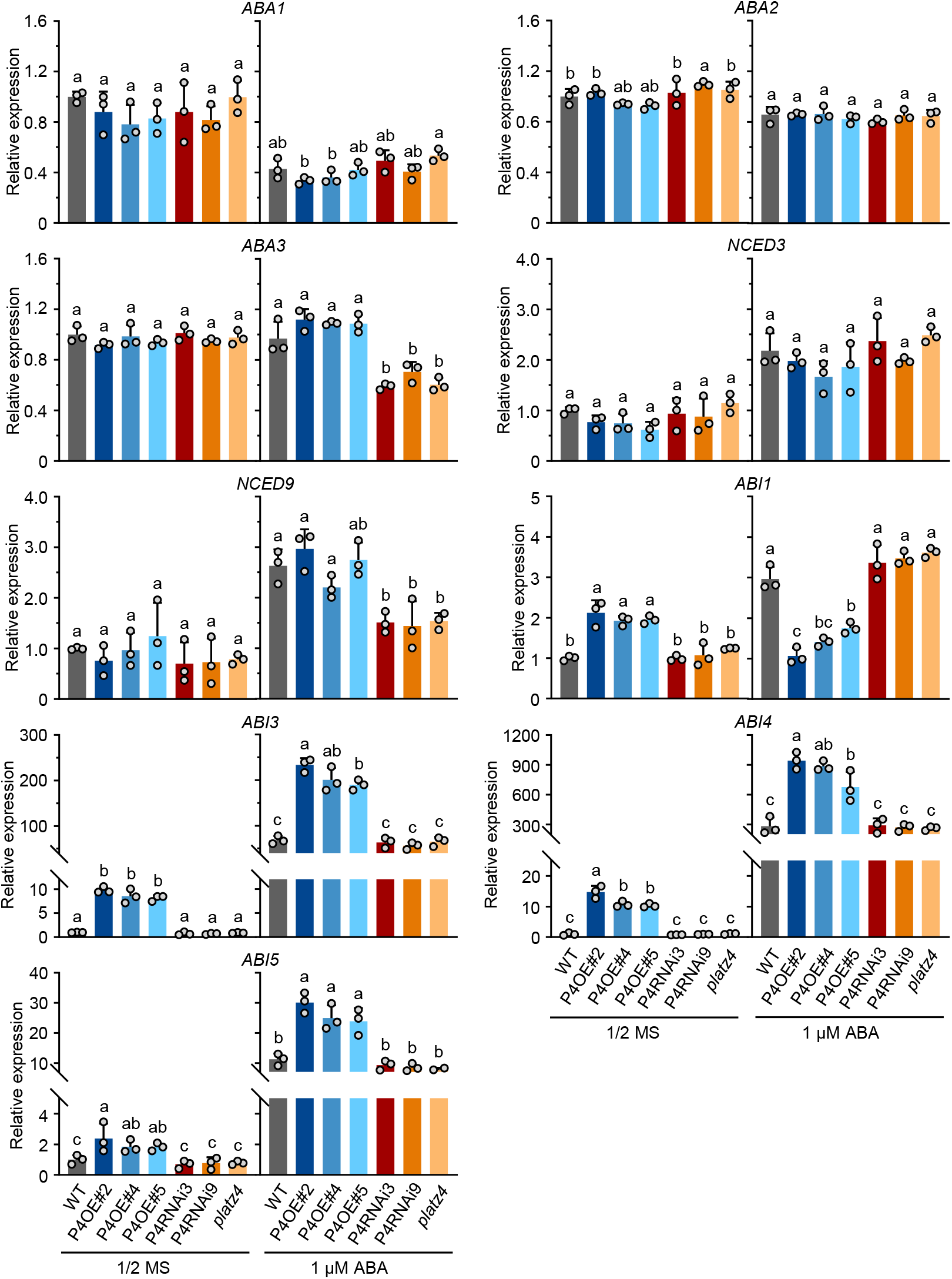
mRNA levels of marker genes involved in ABA biosynthesis and signaling in germinated PLATZ transgenic and WT seeds treated with or without ABA. The expression levels *ABA1, ABA2, ABA3, NCED3, NCED9, ABI1, ABI2, ABI3* and *ABI5* in WT, P4OE#2, P4OE#4, P4OE#5 and P4RNAi3, P4RNAi9, *platz4* germinated seeds grown on half-strength of MS with or without 1 μM ABA for 4 d were analyzed by RT-qPCR. Error bars indicate SEM (*n* = 3). Bars labeled with different lowercase letters are significantly different from one another (*P* < 0.05; one-way ANOVA).

**Supplementary Figure 4.**
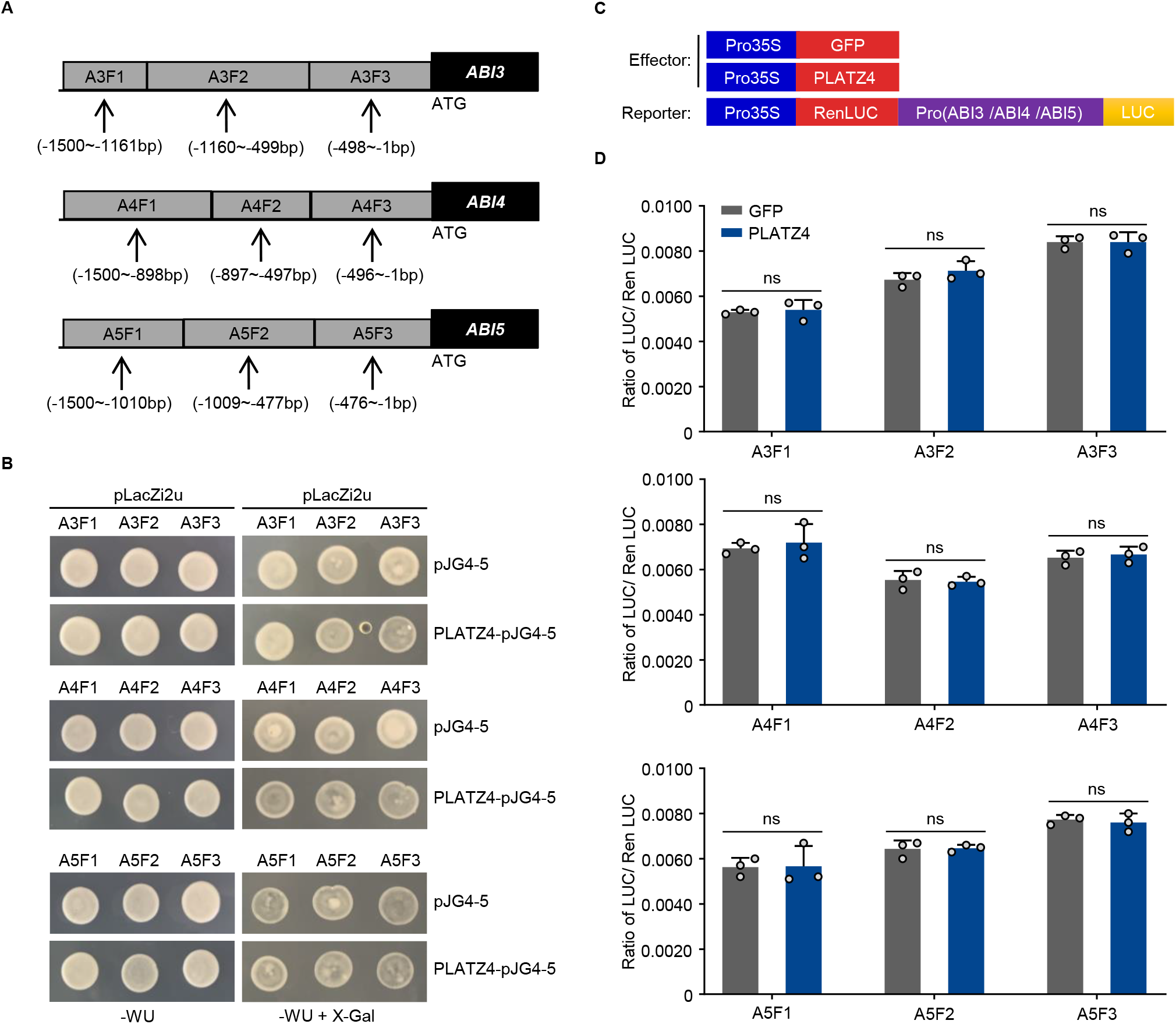
Analysis the binding of PLATZ4 to promoters of *ABI3, ABI4*, and *ABI5*. (A) Diagrams of the promoters and gene regions of *ABI3, ABI4*, and *ABI5*. A3F1–A3F3 indicate the three fragments upstream of *ABI3;* A4F1–A4F3 indicate the three fragments upstream of *ABI4;* and A5F1–A5F3 indicate the three fragments upstream of *ABI5*. (B) Yeast one-hybrid assays revealed the binding of PLATZ4 to promoters of *ABI3, ABI4*, and *ABI5*. Yeast cells were co-transformed with GAL4-Activation Domain (pJG4-5) or PLATZ fused with GAD (PLATZ4-pJG4-5) and the LacZ reporter gene driven by the promoter of *ABI3* (*ProA3F1:LacZ, ProA3F2:LacZ, ProA3F3:LacZ*), *ABI4* (*ProA4F1:LacZ, ProA4F2:LacZ, ProA4F3:LacZ*), and *ABI5* (*ProA5F1:LacZ, ProA5F2:LacZ, ProA5F3:LacZ*), and grown on medium containing X-Gal. Co-expression of GAD and *ABI3* (*ProA3F1:LacZ, ProA3F2:LacZ, ProA3F3:LacZ*), *ABI4* (*ProA4F1:LacZ, ProA4F2:LacZ, ProA4F3:LacZ*), *ABI5* (*ProA5F1:LacZ, ProA5F2:LacZ, ProA5F3:LacZ*) was used as a negative control, respectively. WU, Synthetic Dropout/-Trp-Ura. (C,D) In vivo transient Dual-Luciferase Reporter Assays verified the binding of PLATZ4 to promoters of *ABI3, ABI4, and ABI5* in *N. benthamiana* leaves. The promoters of *ABI3, ABI4, ABI5* were shown in (A). The schematic diagram showed the constructs used in transient expression assays (C), effector constructs contained the PLATZ4 driven by the CaMV35S promoter; reporter constructs contained the three fragments upstream of *ABI3 or ABI4, or ABI5, Pro35S:GFP* was used as a negative control. The values were calculated by the ratio of LUC activities to REN activities (LUC/REN). Error bars indicate SEM (*n* = 3). ns, not significant (Student’s t-test).

**Supplementary Figure 5.**
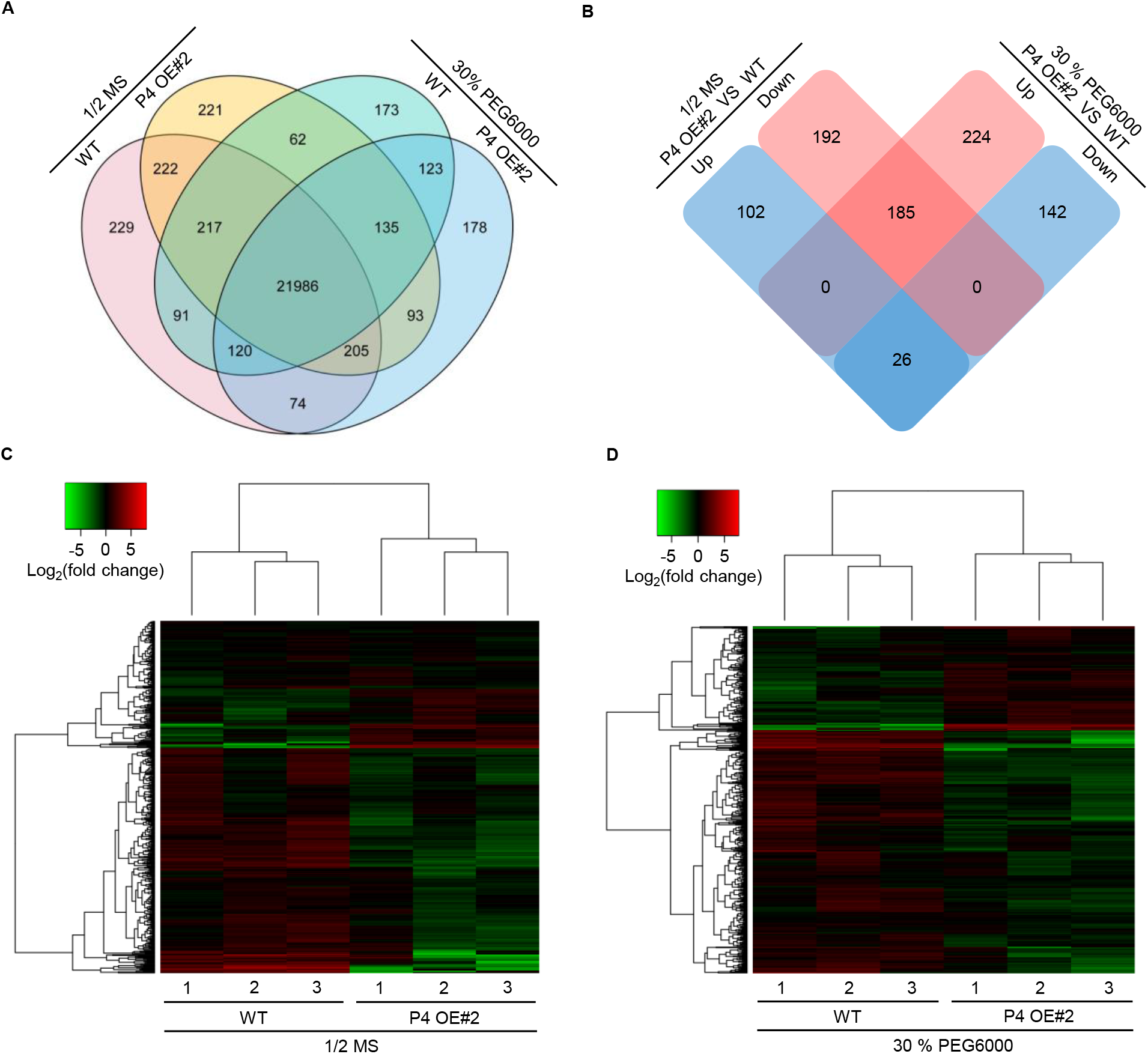
Different expression genes regulated by PLATZ4. (A) Venn diagram of the numbers expressed genes in the P4OE#2 compared with WT before and 3 h after 30 % PEG6000 treatment. (B) Venn diagram of the numbers of differentially expressed genes between P4OE#2 and WT before and after 30 % PEG6000 treatment for 3 h. (C,D) Heatmaps of genes in the P4OE#2 and WT before (C) and 3 h after 30 % PEG6000 treatment (D). In normalized box plots of gene expression: black boxes, no significant difference; red boxes, significant upregulation; green boxes, significant downregulation.

**Supplementary Figure 6.**
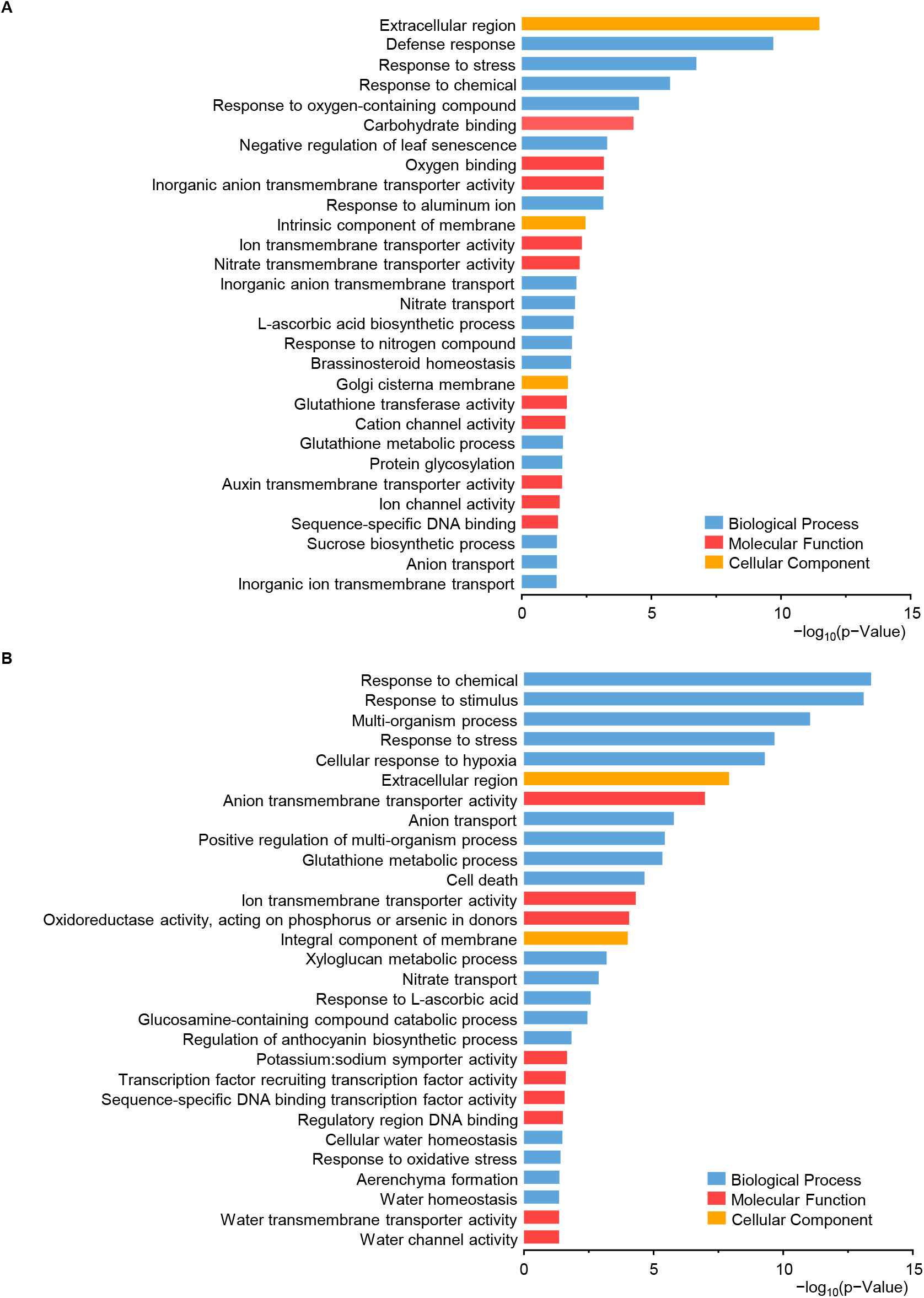
GO analysis of different expression genes regulated by PLATZ4. (A,B) Enrichment analyses of the differentially expressed genes (DEGs) between P4OE#2 and WT before (A) and after 30 % PEG6000 treatment (B) for 3 h, respectively.

**Supplementary Figure 7.**
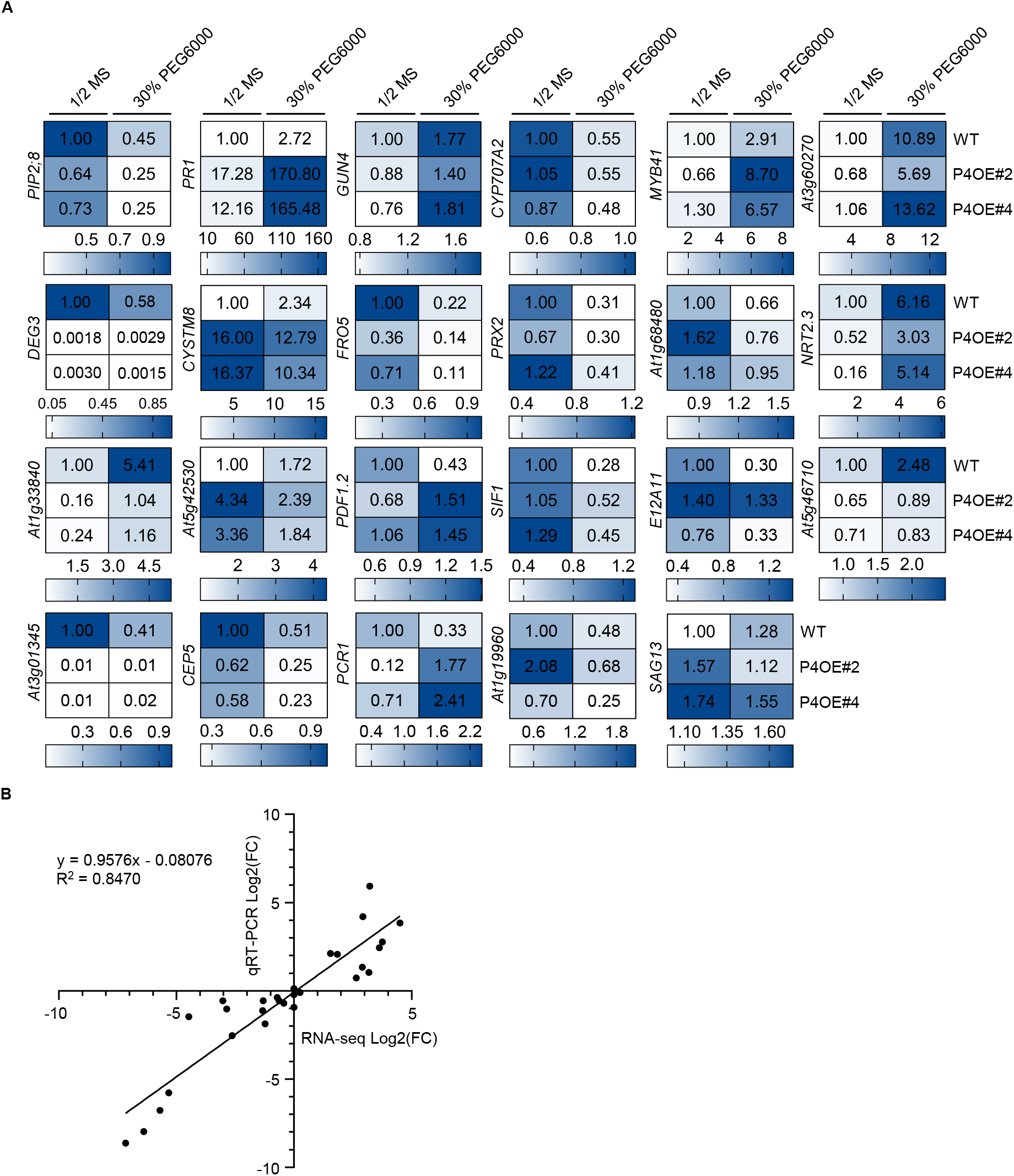
Verification of some DEGs by RT-qPCR. (A) Verification of RNA-seq results by qRT-PCR analyses of 23 representative genes which differentially expressed between P4OE#2, P4OE#4 and WT before and after 30 % PEG6000 treatment for 3 h. Higher transcript levels (OE line/WT) are shown in dark blue and lower transcript levels (OE line/WT) are shown in white. (B) 30 % PEG6000 induction levels of genes determined by qRT-PCR and RNA-seq were closely correlated. FC, fold change.

**Supplementary Figure 8.**
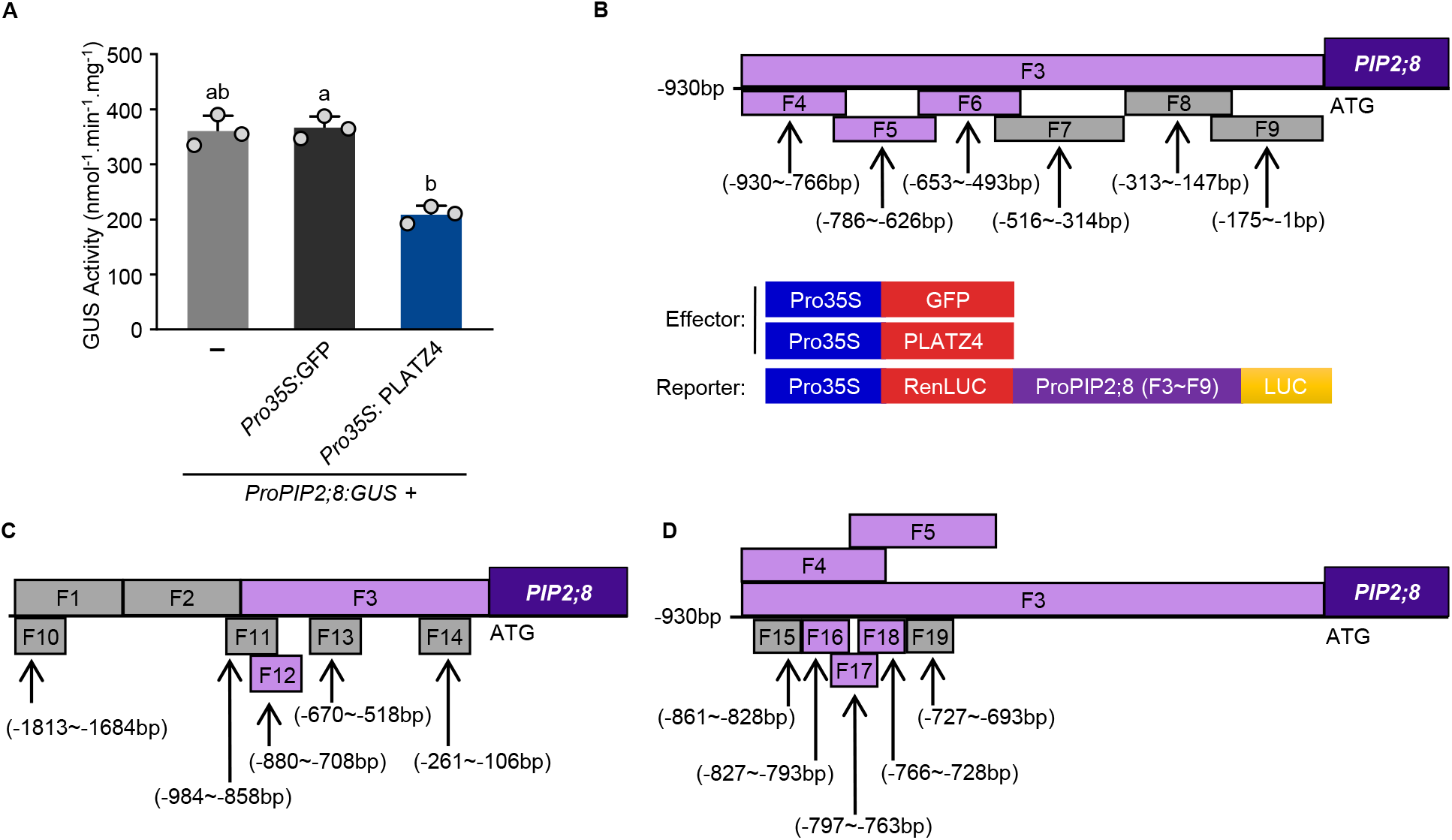
Inhibition of PLATZ4 to *PIP2;8* promoter activity and diagrams of DNA fragments in the promoter of *PIP2;8* in transient Dual-Luciferase Reporter Assay, ChIP-qPCR and EMSAs. (A) PLATZ4 inhibits *PIP2;8* promoter activity. The *ProPIP2;8:GUS* vectors were transformed or co-transformed with the *Pro35S:GFP*, or *Pro35S:PLATZ4* vectors into *N. benthamiana* leaves. The transformed leaves were incubated in darkness for 12 h and maintained in a moist chamber at 25°C for 2 d before the GUS activity was measured. Data represent the mean ±SD of three replicates (*n* = 12). Bars labeled with different lowercase letters are significantly different from one another (P < 0.05; one-way ANOVA). (B) Diagram of DNA fragments in the promoter of *PIP2;8* used in transient Dual-Luciferase Reporter Assays. PLATZ4 driven by the CaMV35S promoter was used as effector constructs. LUC driven by six DNA fragments (F4 to F9) and F3 upstream of *PIP2;8* were used as reporter constructs. And *Pro35S:GFP* was used as a negative control. Gray boxes represented the DNA fragments that PLATZ4 did not bind, while purple boxes represented the DNA fragments that PLATZ4 bound. (C) Diagram of DNA fragments in the promoter of *PIP2;8* used for ChIP-qPCR analysis. F10 to F14 within the F3 DNA fragment were detected. F12 (−880 – −708 bp) DNA fragment was bound by PLATZ4. (D) Diagram of DNA fragments fragments in the promoter of *PIP2;8* used EMSAs. F15 to F19 DNA frangments were detected.

**Supplementary Figure 9.**
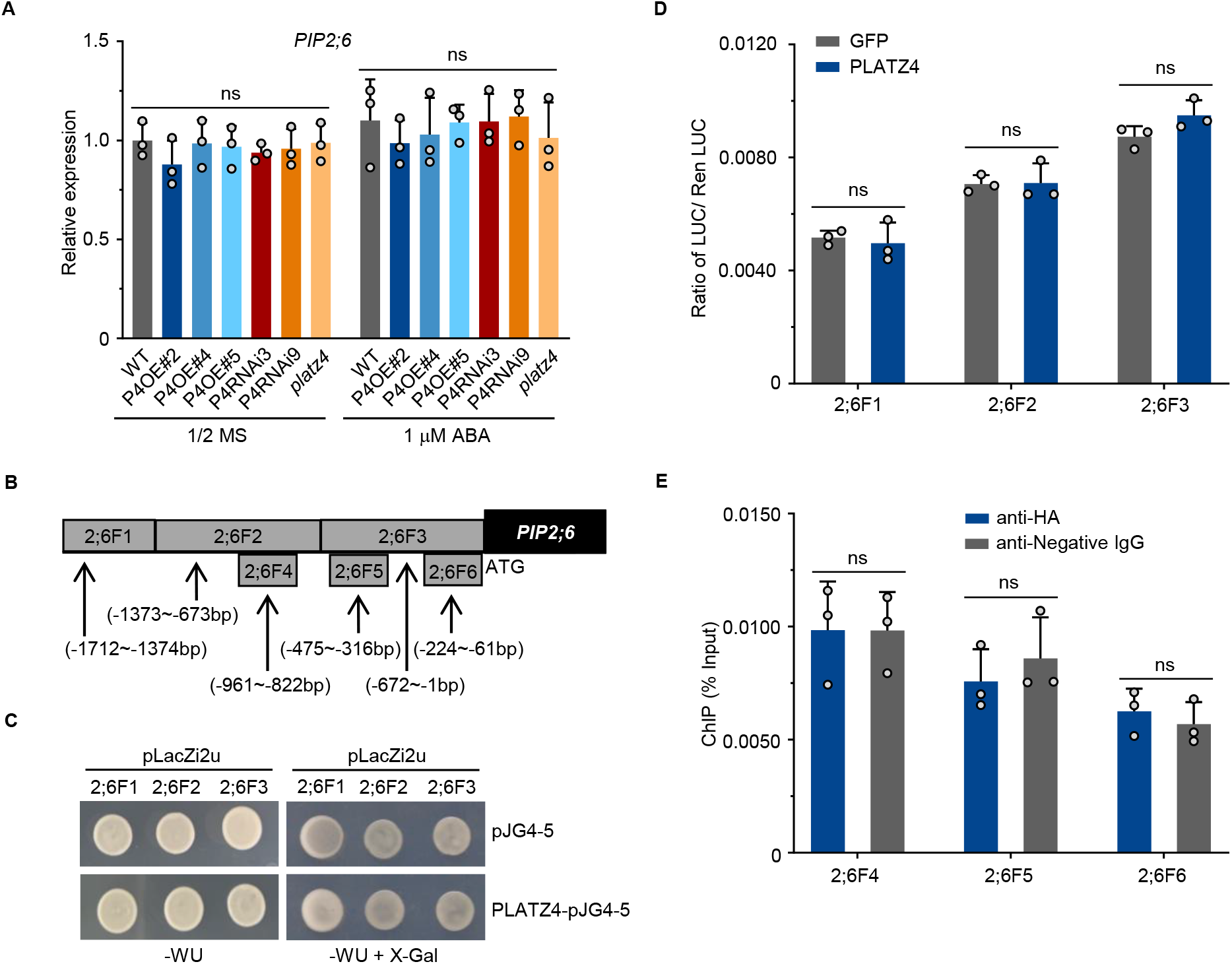
The binding analysis of *PIP2;6* promoter by PLATZ4. (A) The transcript levels of *PIP2;6* in 4-day-old seedlings of WT, P4OE#2, P4OE#4, P4OE#5, P4RNAi3, P4RNAi9, and *platz4* grown under half-strength MS with 30% PEG6000 for 3 h or not. Error bars indicated SEM (*n* = 3). (B) Diagrams of DNA fragments within *PIP2;6* promoter used in Y1H, transient Dual-Luciferase Reporter Assay, and ChIP-qPCR analysis, respectivley. (C) Y1H assays revealed that PLATZ4 did not bind the 2;6F1 to 2;6F3 DNA fragments in the promoter of *PIP2;6*. Yeast cells were co-transformed with GAL4-Activation Domain (pJG4-5) or PLATZ4 fused with GAD (PLATZ4-pJG4-5) and the LacZ reporter gene driven by the indicated DNA fragments (*Pro2;6F1:LacZ, Pro2;6F2:LacZ, Pro2;6F3:LacZ*) and grown on medium containing X-Gal. Co-expression of GAD and *Pro2;6F1:LacZ, Pro2;6F2:LacZ*, or *Pro2;6F3:LacZ* was used as a negative control. WU, Synthetic Dropout/-Trp-Ura. (D) In vivo transient Dual-Luciferase Reporter Assays in *N. benthamiana* leaves. The values were reached by calculating the ratio of LUC activities to REN activities (LUC/REN). Error bars indicate SEM (*n* = 3). ns, not significant (Student’s *t*-test). (E) ChIP-qPCR analysis. Chromatin was isolated from 7-day-old seedlings of the *ProSuper:PLATZ4-HA* transgenic *Arabidopsis* grown on half-strength of MS medium. Immunoprecipitation was carried out with the HA antibody. Then the F4 to F6 DNA fragments in the *PIP2;6* promoter were quantified using PCR. Error bars indicate SEM (*n* = 3). ns, not significant (Student’s t-test).

**Supplementary Figure 10.**
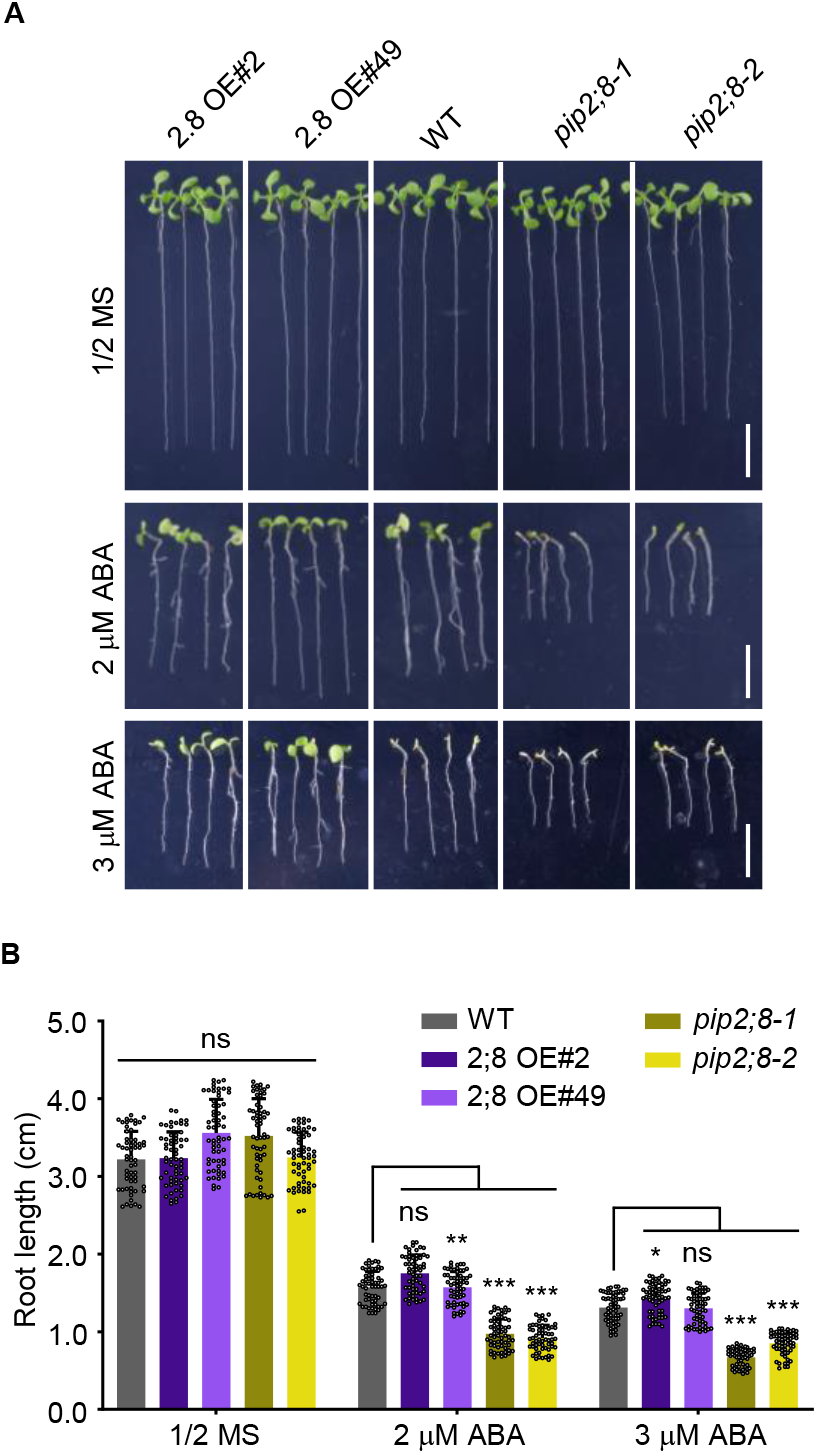
PIP2;8 negatively regulates ABA response in Arabidopsis seedlings. (A) Phenotypes of WT, 2;8OE#2, 2;8OE#49 and *pip2;8-1, pip2;8-2* plants grown on half-strength MS medium with or without 2 or 3 μM ABA for 8 d. (B) The primary root length of the seedlings shown in (A). Error bars indicated SEM (*n* = 60). The asterisks indicate significant differences compared with WT. ns, not significant; *, *P* < 0.05; **, *P* < 0.01; ***, *P* < 0.001 (oneway ANOVA). These experiments were repeated three times with similar results.

**Supplementary Figure 11.**
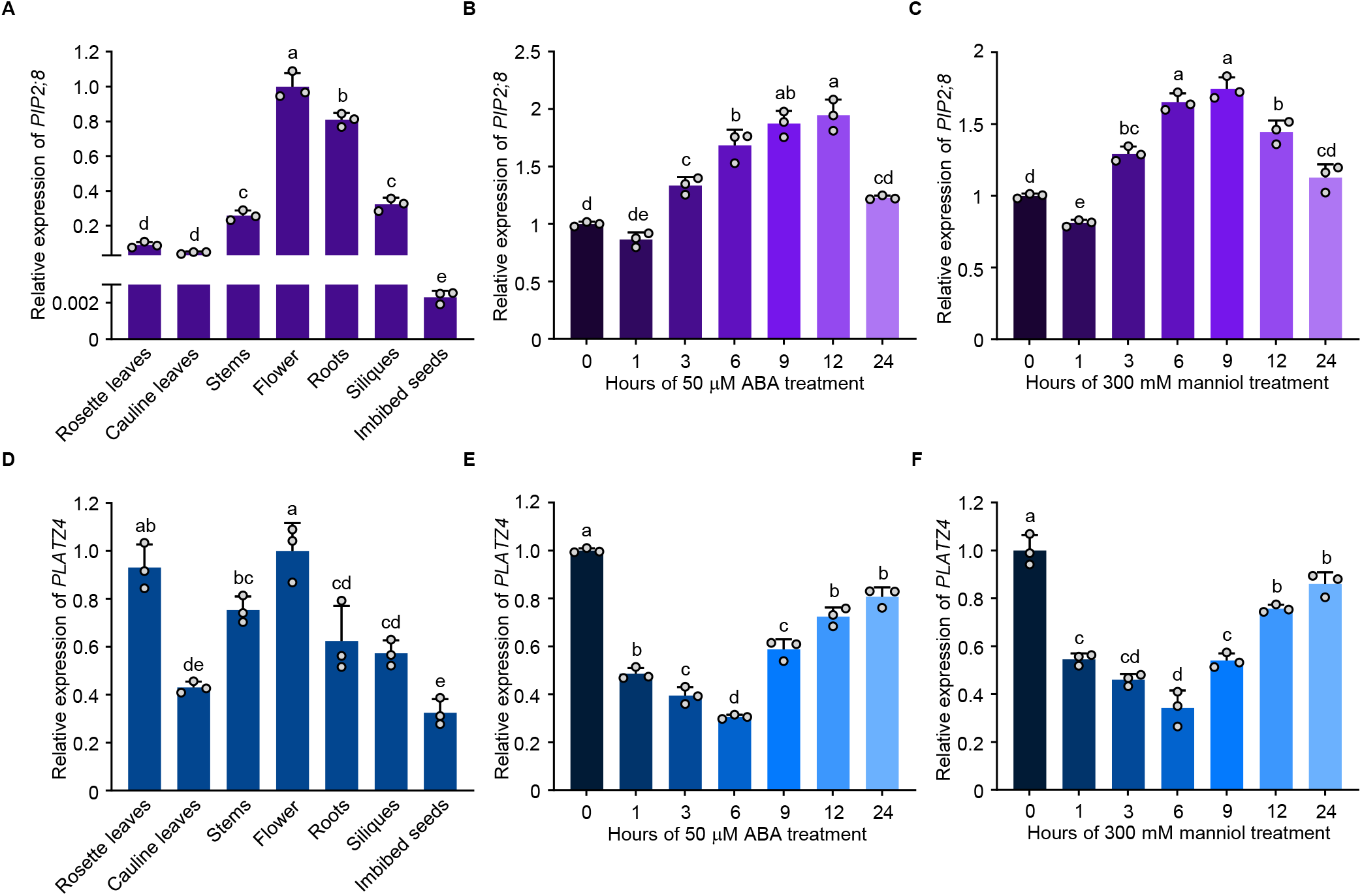
Expression patterns of *PIP2;8* and *PLATZ4*. (A) Expression analysis of *PIP2;8* in roots, stems, rosette leaves, cauline leaves, flower, siliques, and seeds by RT–qPCR. Data were normalized against the expression of *GAPDH* and *UBQ10*. Error bars indicate SEM (*n* = 3). Bars labeled with different lowercase letters are significantly different from one another (*P* < 0.05; one-way ANOVA). (B,C) Responsiveness of *PIP2;8* in the 10-day-old seedlings treated with 50 μM ABA, or 300 mM mannitol for 0, 1, 3, 6, 9, 12, and 24 h. Data were normalized against the expression of *GAPDH* and *UBQ10*. The means were calculated from three independent replicates and compared with the no-treatment condition (0 h). Error bars indicate SEM (*n* = 3). Bars labeled with different lowercase letters are significantly different from one another (*P* < 0.05; one-way ANOVA). (D) Expression analysis of *PLATZ4* in roots, stems, rosette leaves, cauline leaves, flowers, siliques, and seeds by RT–qPCR. Data were normalized against the expression of *GAPDH* and *UBQ10*. Error bars indicate SEM (*n* = 3). Bars labeled with different lowercase letters are significantly different from one another (*P* < 0.05; one-way ANOVA). (E,F) Responsiveness of *PLATZ4* in the 10-day-old seedlings treated with 50 μM ABA, or 300 mM mannitol for 0, 1, 3, 6, 9, 12, and 24 h. Data were normalized against the expression of *GAPDH* and *UBQ10*. The means were calculated from three independent replicates and compared with the no-treatment condition (0 h). Error bars indicate SEM (*n* = 3). Bars labeled with different lowercase letters are significantly different from one another (*P* < 0.05; one-way ANOVA).

**Supplementary Figure 12.**
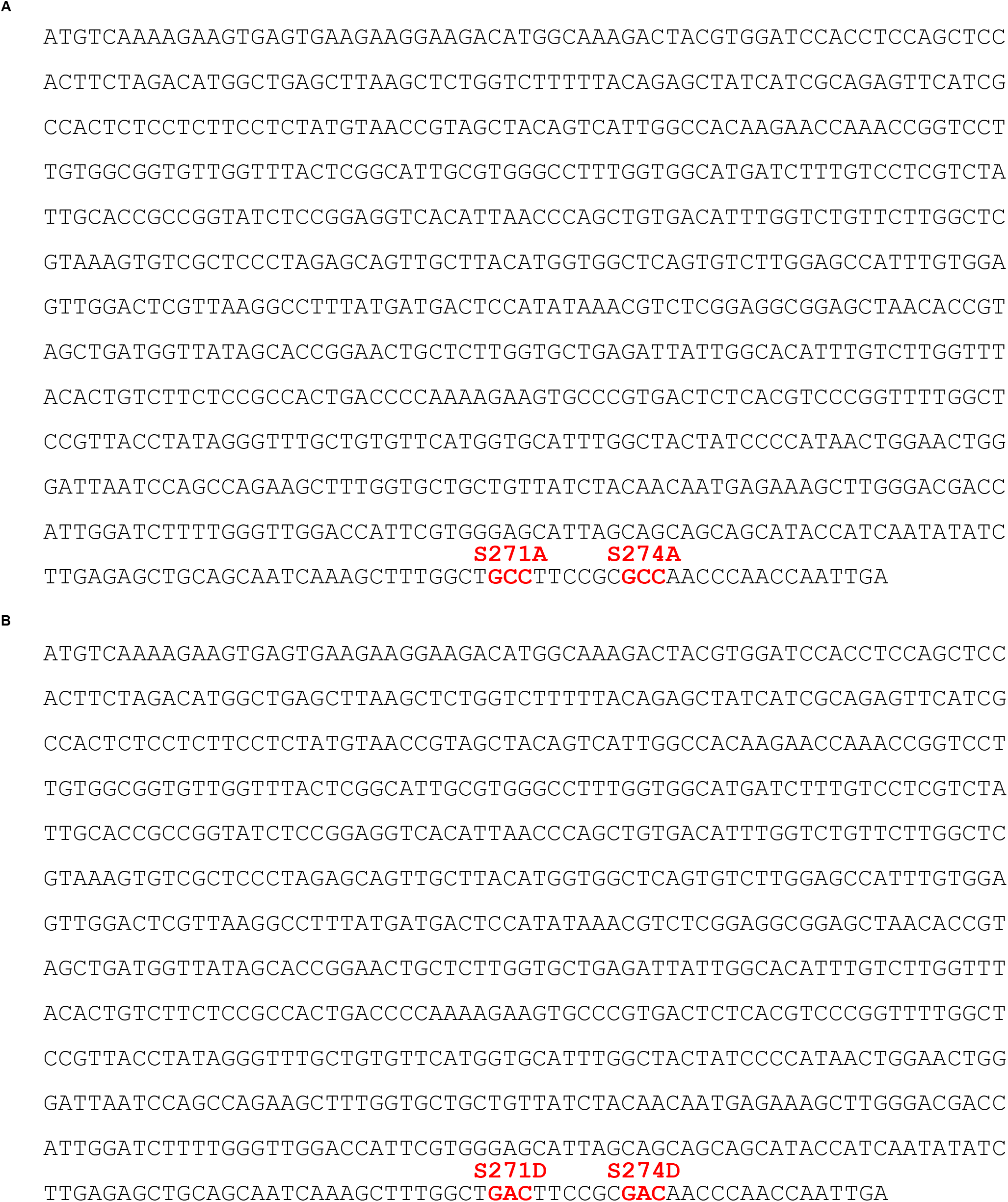
Site-directed mutagenesis of the *PIP2;8* sequence. (A) Candidate phosphorylation site (S271 and S274) is mutated to alanine (S271A and S271A) respectively, which is highlighted in red in the cDNA sequence of *PIP2;8*. (B) Candidate phosphorylation site (S271 and S274) is mutated to aspartic acid (S271D and S271D) respectively, which is highlighted in red in the DNA sequence of *PIP2;8*.

**Supplementary Figure 13.**
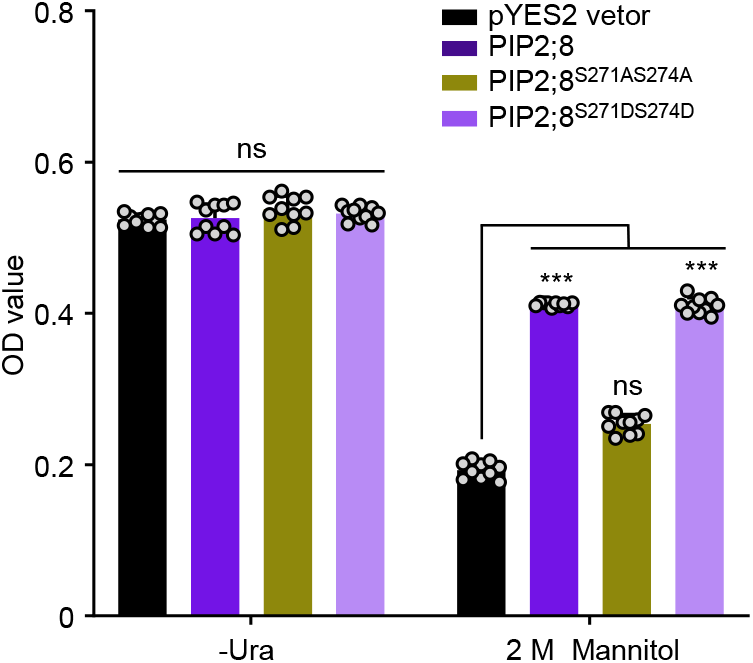
Effects of overexpression of *PIP2;8, PIP2;8^S271DS274D^, PIP2;8^S271AS274A^* on the growth of yeast cells under osmotic stress. Yeast cells were transformed *PIP2;8, PIP2;8^S271DS274D^, PIP2;8^S271AS274A^* and grown under 2 M Mannitol mannitol at 30 °C for 36 h or not. Yeast cells transformed with pYES2 empty vector were used as negative control. Optical densities (OD) at 600 nm of the independent transformants were recorded. Error bars indicate SEM (*n* = 10). ns, not significant; ***, *P* < 0.001 (one-way ANOVA).

**Supplementary Figure 14.**
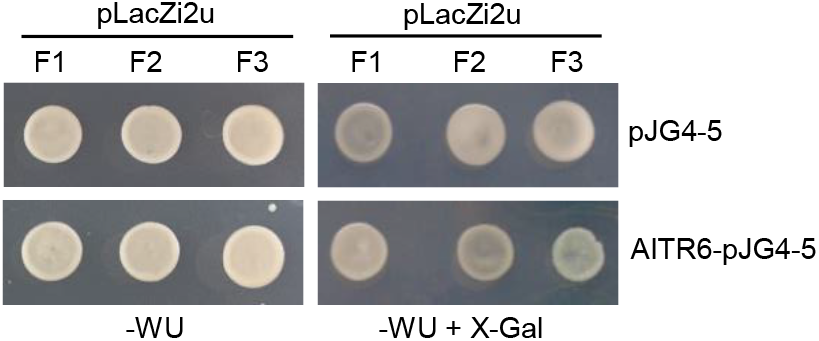
The binding analysis of *PIP2;8* promoter by AITR6 by Y1H. Yeast cells were co-transformed with GAL4-Activation Domain (pJG4-5) or AITR6 fused with GAD (AITR6-pJG4-5), and the LacZ reporter gene driven by the promoter of PIP2;8 (*ProF1:LacZ, ProF2:LacZ, ProF3:LacZ*) and grown on medium containing X-Gal. Co-expression of pJG4-5 and *ProF1:LacZ, ProF2: LacZ*, or *ProF3:LacZ* were used as a negative control. WU, Synthetic Dropout/-Trp-Ura.

